# Fractions strategy differences in those born extremely preterm

**DOI:** 10.1101/2022.12.06.519175

**Authors:** Sarah Carr, W. Michael Babinchak, Ana Istrate, Blaine Martyn-Dow, George Wang, Weicong Chen, Jeremy Fondran, Jing Zhang, Michael Wien, Seo Yeon Yoon, Anne Birnbaum, Elizabeth Roth, Carol Gross, Nori Minich, Lee Thompson, Won Hwa Kim, Yaakov Stern, Chiara Nosarti, H. Gerry Taylor, Curtis Tatsuoka

## Abstract

**Introduction:** To investigate the effects of different strategies and cognitive load we explored brain hemodynamic responses associated with the use of different strategies to solve subtraction of fractions. We focused on those born extremely preterm (EPT; <28 weeks’ gestation) as they are known to have cognitive challenges and struggle with mathematics. We also included a group of full-term (FT) peers for comparison.

**Methods:** Functional MRI was acquired while the participants mentally solved fraction equations using either a strategy based on improper or mixed fractions. Different fraction item types were given, which affected respective required cognitive loads per strategy. Diffusion and T1-weighted structural images were also acquired.

**Results:** The EPT and FT groups differed in terms of task-related hemodynamic responses. Functional group differences were greatest when strategies were applied to item types that result in high cognitive load. Other findings showed reduced white and grey matter volume and reduced white matter connectivity in widespread areas in the EPT group compared to the FT group.

**Conclusion:** The understanding of function and structure presented here may help inform pedagogical practices by allowing for tailoring of mathematical education through identifying suitable strategy adoption that depends on item type, to circumvent weaknesses in cognitive skills.

## 1.0 Introduction

Extremely preterm (EPT) birth is defined as birth before 28 completed weeks of gestation. The incidence of preterm birth (gestational age (GA) < 37 weeks) in the US is about one in ten births [1] and globally, 2-5 births in 1000 are EPT [2]. Survival rates among those born EPT are around 33% at 23 weeks’ GA, 65% at 24 weeks, 81% at 25 weeks and 94% at 27 weeks [3]. Children born EPT often have deficits in working memory, executive function, inhibition, and processing speed when compared to full-term (FT) control subjects [4, 5]. Children born EPT are known to have cognitive difficulties through adolescence and by early adulthood they are likely to have lower IQ and impaired visuomotor, prospective memory, language and executive functioning skills [6]. Therefore, EPT individuals are vulnerable to experience impairments in a range of neurocognitive skills that are crucial for learning, problem solving and effective communication, which are likely to exert a negative impact on their overall academic achievement [7]. Mathematics difficulties have been associated with poor performance on working memory and visuo-spatial processing tasks, rather than impairments in numerical representation [4], suggesting that the nature of mathematics difficulties of preterm children is different from that of individuals with developmental dyscalculia.

In FT individuals, multiple brain regions have been implicated in the neurobiology of computation, including in the frontal, parietal, and temporal cortices, as well as subcortically, in the thalamus, basal ganglia, and hippocampus [8, 9]. The intraparietal sulcus (IPS) appears to be the primary locus of numerical processing, and that it is critical to mathematical reasoning but not necessarily specific [10]. Nevertheless, the IPS is considered as part of a “core” visual-spatial number system, along with the fusiform gyrus [10, 11]. Tract-specific white matter microstructural characteristics, such as fractional anisotropy (FA), have been associated with mathematical skill. These tracks are the superior longitudinal fasciculus (connecting fronto-parietal cortices) [12-14], inferior longitudinal fasciculus (fronto-occipito-temporal areas) [12, 15] and arcurate fasciculus (inferior fronto-occipito-temporal, connecting Wernicke’s and Broca’s areas) [16]. The processing and maintenance of both visual and verbal information is also often central to mathematical performance [17-19].

Few functional imaging studies have investigated the neural correlates associated with performing mathematics in preterm-born individuals. Functional magnetic resonance imaging (fMRI) studies, involving tasks such as the Stroop and number estimation, have shown an association between longer gestation and more pronounced hemodynamic responses in parietal brain regions. Shorter GA has also been associated with more prefrontal cortex activity in young adults born < 37 weeks’ gestation when completing a magnitude comparison task [20], in 6- to 7-year-old children born between 27-32 weeks with a numerical distance effect task [21, 22] and 11-year-old youth born < 28 weeks’ gestation with a working memory/selective attention task [23]. As FT children mature, areas of the brain associated with performing mathematics change from mostly frontal to less frontal and more parietal areas, possibly due to procedural changes in how mathematics is performed [24, 25]. Individuals born EPT may thus have a more ‘immature’ mathematical network [20].

A study of functional connectivity and performance on mathematical tests showed patterns of association unique to preterm individuals born between 25-36 weeks’ gestation. Functional connectivity derived from resting state fMRI in adulthood (26 years old) showed associations with mathematical ability in childhood in bilateral fronto-parietal networks in preterm adults that were different to FT adults [26]. Also, mathematical abilities recorded in childhood (8 years old) were more positively associated with adult IQ for preterm than for FT individuals.

A study of mathematics and regional brain volumes found an association between reduced grey matter volume in left parietal lobe and poor mathematics performance in EPT teenagers [27]. Diffusion weighted imaging studies assessing white matter characteristics have shown that EPT infants scanned at 38 weeks’ post-conceptual age have alterations in white matter microstructure (reduced FA and increased radial diffusivity) in widespread areas associated with early myelination (such as sensorimotor areas) [28] and frontal and occipital areas [29] compared with FT babies. In older EPT children, 9-16 years of age, lower FA values in the corpus callosum, forceps minor and major, cingulate gyrus, inferior fronto-occipital fasciculus, inferior and superior longitudinal fasciculi and corticospinal tract have been associated with impaired executive functioning [30]. In EPT infants scanned between birth and 4 years of age, changes in mean diffusivity values in the internal and external capsules have been associated with intelligence and language [31]. The long-range connections between frontal, temporal and parietal lobes serve to integrate and control complex brain processes involved in cognition and skills. The authors are not aware of any research investigating mathematics and white matter connectivity in individuals who were born EPT. In summary, the EPT population has a multitude of structural and functional brain changes that affect both their ability to do mathematics and domain-general functions such as attention and working memory.

The present study represents an initial effort to identify individual differences in brain structure and function as a means for developing more targeted approaches to mathematical instruction. We focus on fractions, as this mathematics domain poses demands on multiple computational abilities, requires more advanced skills than arithmetic problem-solving, and is a key mathematical subject areaWe focused on adolescents as they adapt well as an age group to fMRI experimentation and are mathematically advanced enough to have already had instruction in solving fraction problems. The goal was to investigate group differences in hemodynamic response patterns during processing of same types of test items invoking different strategies that varied in cognitive load. We also aimed to explore between group differences in brain structure. We hypothesized that there would be differences between FT and EPT groups in hemodynamic response patterns, particularly for the math problem and strategy combinations that require relatively higher cognitive loads. Discovery of group differences could suggest the need for stronger emphasis on flexible strategy adoption for adolescents born EPT that would allow for reductions in cognitive load.

## 2.0 Methods

### 2.1 Participants

The participants made two visits to University Hospitals Cleveland Medical Center (UHCMC) on separate days. The first visit was to undertake neuropsychological testing and training in fractions, (lasting 3 - 4 hours). The second visit was to the MRI department, lasting around 2 hours. Duration between visits ranged from 7 to 364 days with the median duration being 22 days (SD 81.2 days). In total 50 subjects attended for neuropsychological training and a subset was selected for MRI based on: 1) the subject’s willingness to participate in an MRI, 2) the lack of physical contraindications to MRI (e.g., claustrophobia, the presence of cardiac pacemakers or other medical devices, metallic fragments such as shrapnel, etc.), and 3) competency in doing fractions without pen and paper. A total of 30 subjects underwent MRI, including 14 EPT adolescents, all with either GA <28 weeks and/or birth weight < 1000 g. EPT participants were 15-17 years of age and included 12 females. The 16 FT adolescences were aged 15-17 years old and included 8 males. The FT youth were recruited from the same regular classrooms as the children born extremely preterm and were thus representative of children attending regular education classes. Eleven EPT and 15 FT participants were right-handed, 2 EPT and 1 FT participants were left-handed and 1 EPT participant was ambidextrous. Participant characteristics are summarized in Table 1.

**Table 1:**
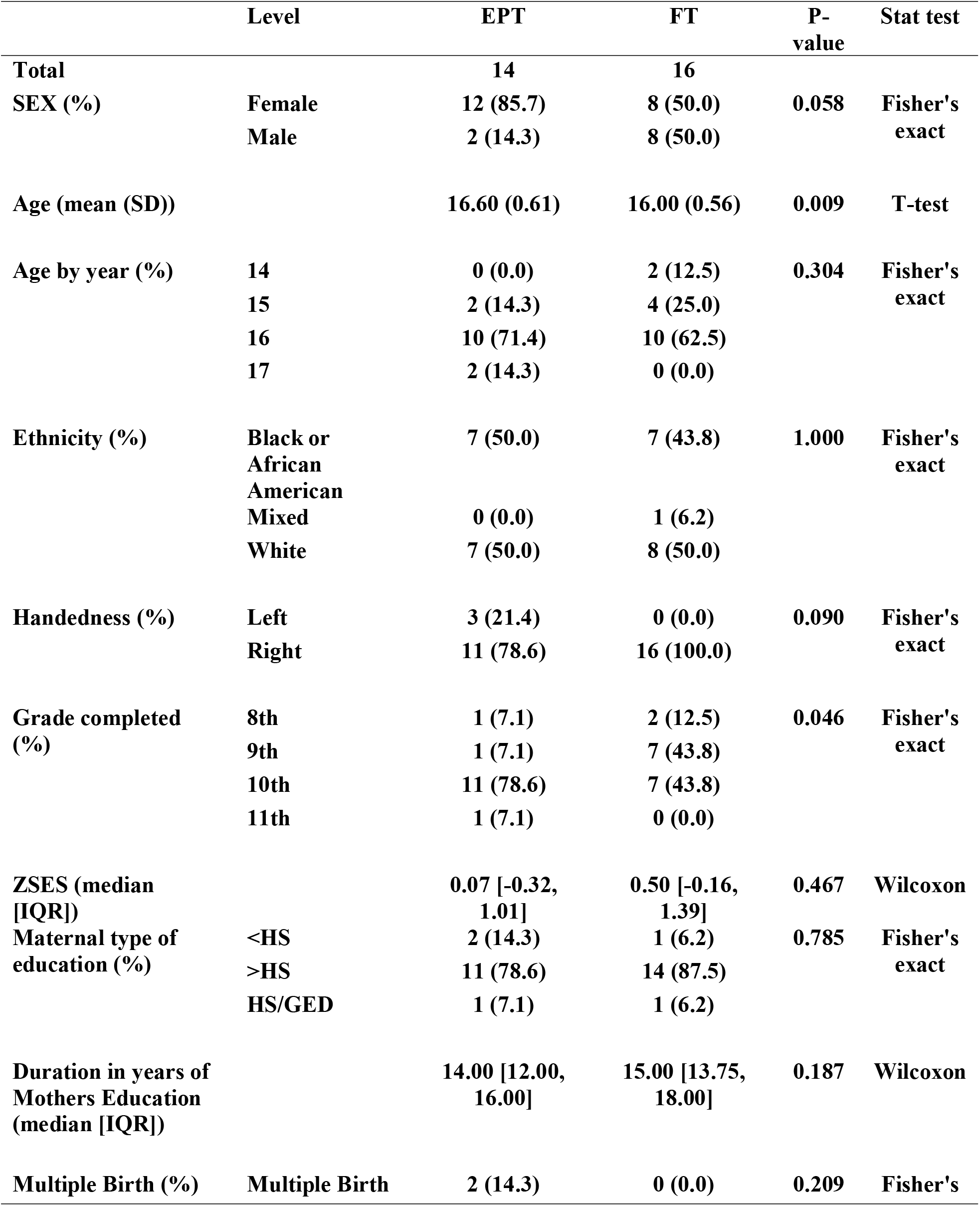
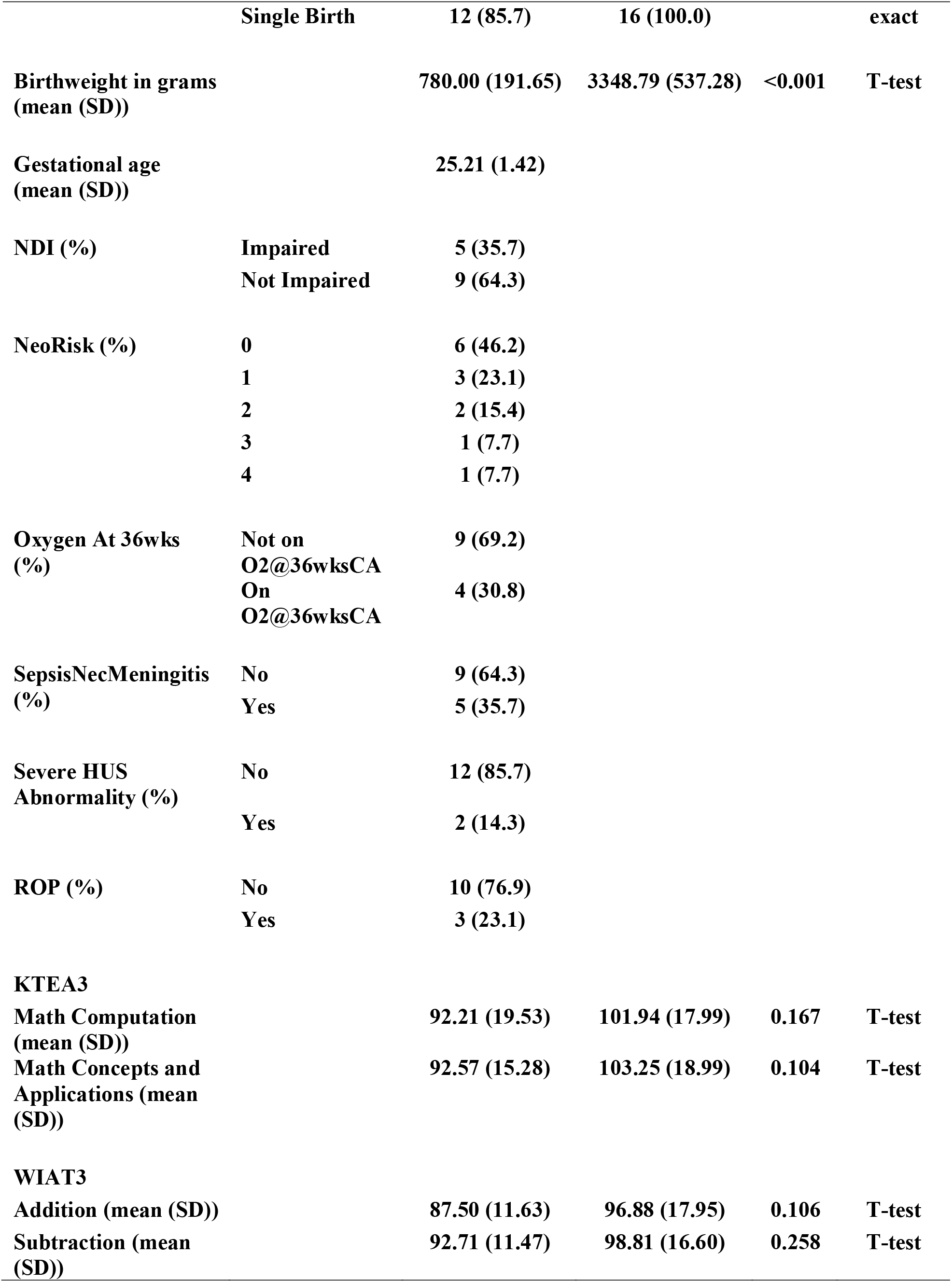
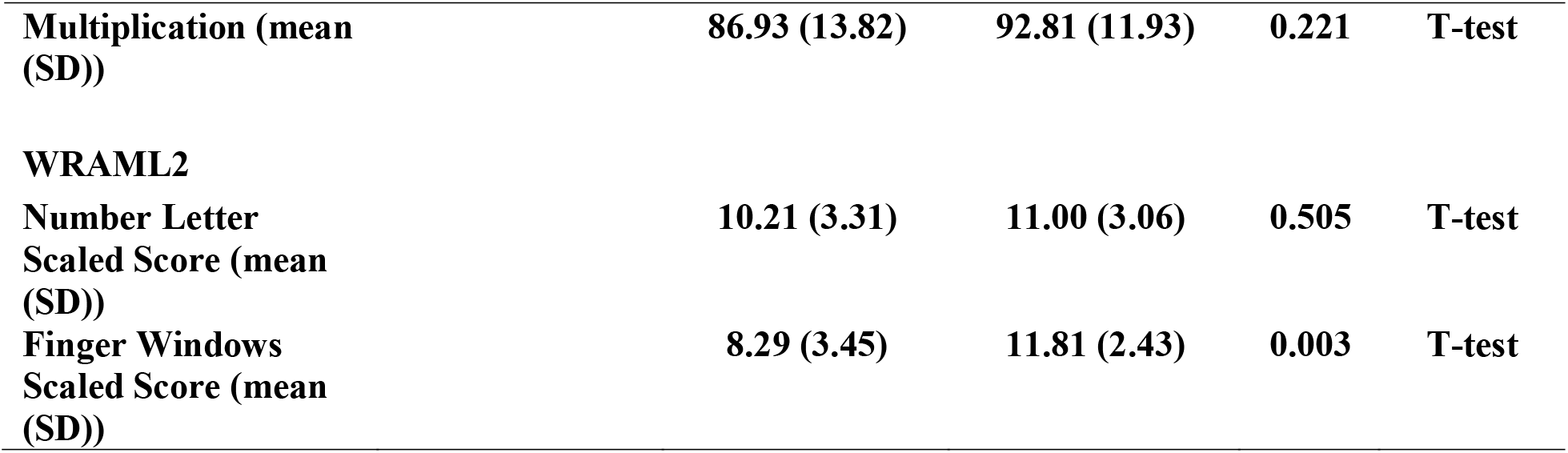
Summary of all subjects recruited for neuropsychological assessment. A subset of 30 subjects received MRI. Normally distributed continuous variables are described using mean (SD) and tested using a t-test. Not normally distributed continuous variables are described using median [IQR, interquartile range] and tested using a Wilcoxon test. Categorical variables are described using percentages and tested using the Chi-squared test or Fisher’s exact test (for variables with small values). SD = Standard deviation, ZSES = a composite (mean of sample z scores) of maternal education level, a measure of social advantage based on parent occupation, and medium income of the family. HS = high school, GED = general education degree certificate, NDI = neurodevelopment impairment (CP, Hearing, Vision and/or MDI<70), NeoRisk = number of neonatal risk factors, SepsisNecMeningitis = sepsis, necrotising enterocolitis or menigitis present, HUS = Head UltraSound, ROP = Retinopathy of prematurity, KTEA3 = Kaufman Test of Educations Achievement third edition, WIAT3 = Wechsler Individual Achievement Test third edition.WRAML2 = Wide Range Assessment of Memory and Learning, second edition.

The Institutional Review Board office at UHCMC granted ethics approval prior to the study. The ethical considerations complied with the Declaration of Helsinki for human subject research. Subjects gave informed assent and their parents gave informed consent prior to taking part.

### 2.2 Neuropsychological testing

Neuropsychological testing was performed using the NIH toolbox - https://www.healthmeasures.net/explore-measurement-systems/nih-toolbox. It includes a range of tests to assess cognition, emotion, sensory and motor function. The cognition tests assessed executive function, working memory, verbal fluency, attention, processing speed and episodic memory. In the interests of brevity, the reader is directed to [32] for full details of the tests. Test score summaries of working memory as assessed by WRAML2 (Wide Range Assessment of Memory and Learning, second edition), by birth group, are also reported in Table 1.

### 2.3 Fractions strategies

We have developed a set of problems assessing subtraction of mixed fractions for administration in the scanner that are similar to a diagnostic subtraction of mixed fractions test developed by K. Tatsuoka [33, 34]. This test and its item-skill specifications have been widely studied and validated, with data available at the Royal Statistical Society website (https://rss.org.uk) [35]. The problems are solved using mental arithmetic. Each item can be solved with one of two strategies, with the required strategy prompted during the fMRI session. Item types are described as:

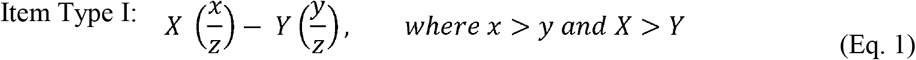

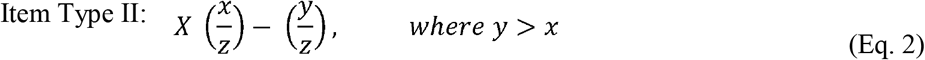

Solving strategies include “mixed” and “improper” methods. Consider 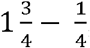, an Item Type I problem. The “mixed” method involves separate computations on whole number and fraction components, e.g.,

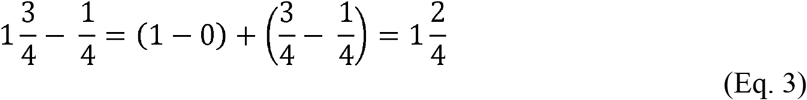

while the “improper” strategy involves solving using improper fractions by converting a whole number to fraction, e.g.,

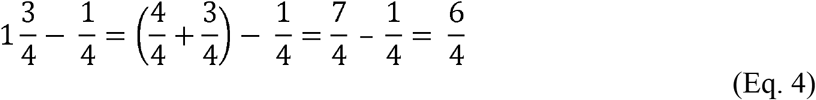

Note that in solving an item of this type, the improper strategy requires higher cognitive load, specifically in working memory. However, the relative cognitive demands or difficulty levels imposed by the two strategies (easier vs. harder) vary by the type of problem. For an Item Type I problem, the mixed method requires fewer steps, and thus a lower cognitive load, than does the improper fraction method. However, for an Item Type II problem, the improper fraction approach is more efficient than the mixed fraction method. A problem of this type would be: 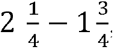, which is an Item Type II.

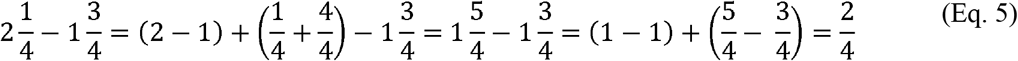

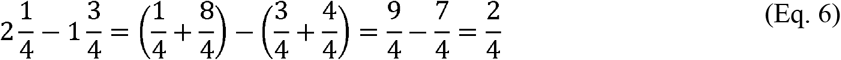

In this problem the mixed fractions strategy requires borrowing from a whole number, i.e. 1 or 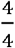 from the 2 on the left-hand side of the problem, yielding 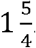. In this case, and more generally for items of Type II, converting both numbers to improper fractions before subtracting places similar or even fewer demands on working memory. This can be made more explicit by listing out the computations and steps and numbers that must be held in memory during the problem-solving process. In general, cognitive load varies depending on strategy and item type. Depending on item type, one strategy can impose a higher demand on working memory than the other. See Figure 1.

**Figure 1:**
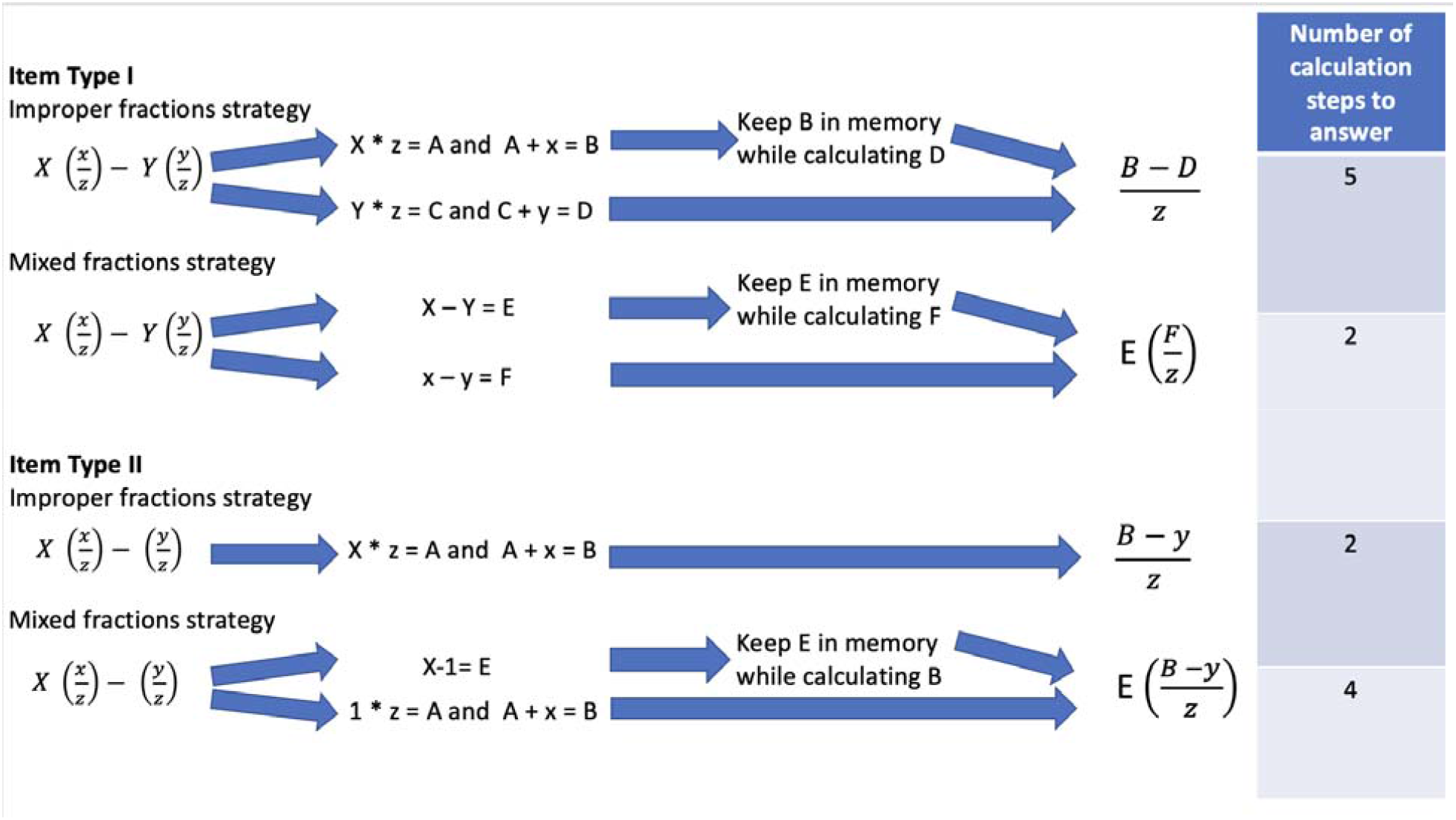
Strategy/Item Type calculation steps demonstrating the cognitive load for each combination.

For the two item types that we considered for fMRI, there are thus harder and easier difficulty levels for each of the two strategies. For the items considered, when one strategy is at the harder level, the other strategy is at the easier level, as illustrated above. This suggests that more difficult levels of specific strategies can be avoided through flexibility in adopting alternative strategies. The present study examines associations of different problem-solving strategies that have been taught and practiced to mastery by the participants with patterns of neural activation on fMRI.

### 2.4 Fraction Strategy Training

Training in subtraction of mixed fractions was carried out as part of the initial visit, through a multi-step program that began with review of basic concepts followed by instruction and practice with the two solving strategies. See Supplement section S1.1 for further details. The majority of subjects from both the EPT and FT groups were able to achieve a high level of accuracy on the final computer administration training problems, median values for EPT - 32.5 out of 35 (IQR 4.5) and FT - 34 out of 35 (IQR 1.0).

### 2.5 fMRI Stimulus protocols

Respective blocks of items requiring the two fraction solving strategies were presented separately in the scanner. An example sequence is shown in Figure 2. Which strategy to use was prompted through instructions printed on a screen at the beginning of each scan and with the presentation of each equation. Colored *X*s were used to prompt the format of the expected solution, e.g.:

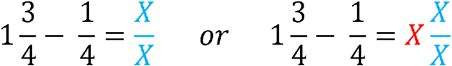

**Figure 2:**
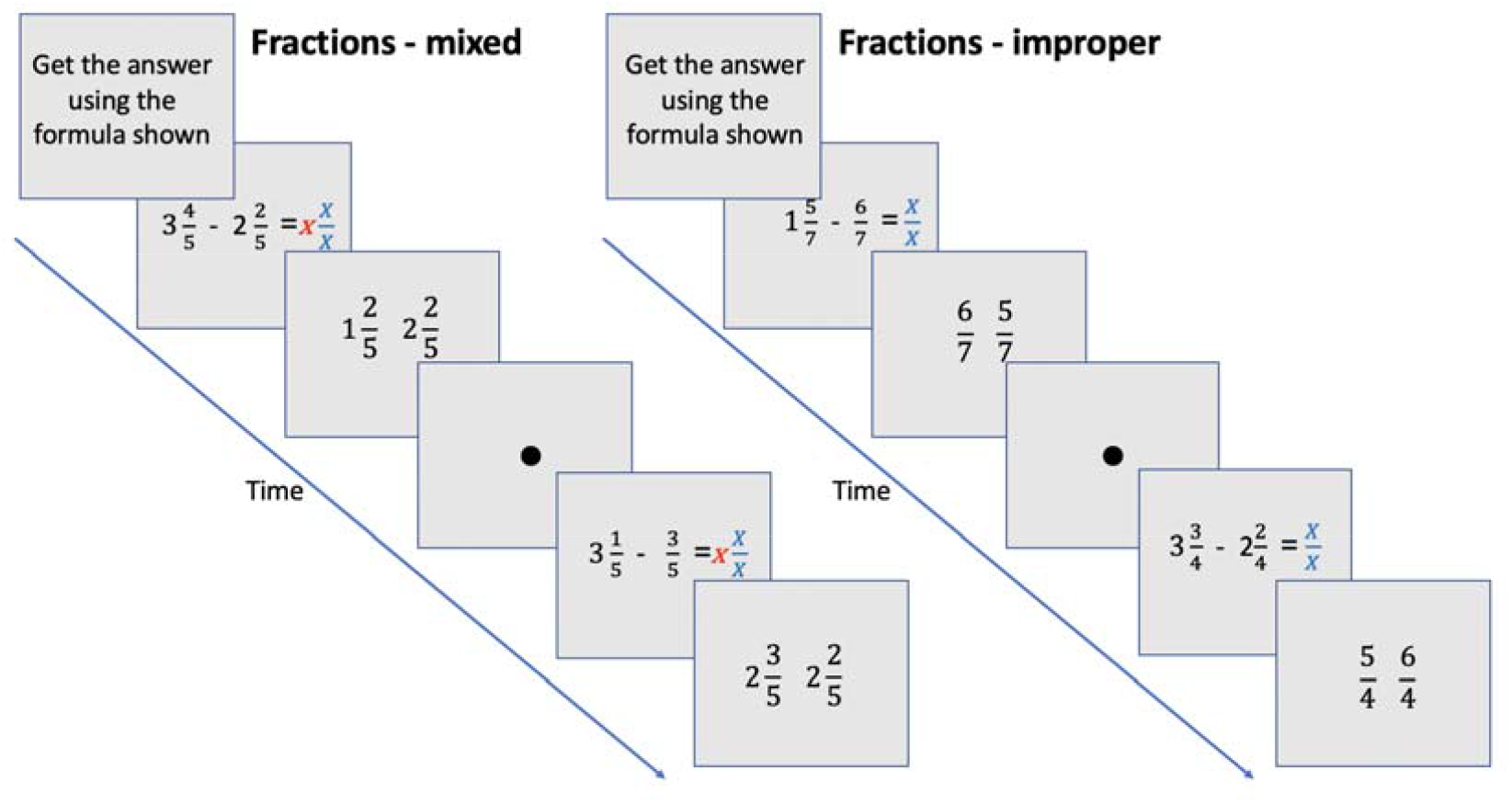
Sample fraction protocol demonstrating the two strategies.

Correct strategy adoption was validated by high correctness rates. See Figure 2 for further details. For each strategy, Item Types I and II were presented within each scan in an interleaved fashion. For the improper fractions strategy, the Item Type II involves 3 steps, while Item Type I involves 5 steps. For mixed fractions, Item Type I is only 2 steps, while Item Type II is 4 steps. A fraction was displayed on the screen for 12-18 seconds, depending on difficulty level. Difficulty was determined by the cognitive load/number of steps required to solve the equation. In the response phase of solving an item, another screen was then displayed with two possible answers to choose from. The answers were displayed in the format related to the strategy in use, and displayed for only 6 seconds. There was thus insufficient time for subjects to convert their answer if they had used the incorrect strategy, which encouraged correct adoption. Subjects indicated their answer choice by pressing a button on a response box held in the right hand. Finally, the presentation of each fraction item was separated by a ‘rest’ condition. A black dot was displayed in the center of the screen and subjects were instructed before the scan to look at the dot for the duration. See Supplement section S1.2 for further information about the presentation of the stimulus.

### 2.6 MR Imaging

Scans were acquired using a Philips Ingenuity 3T PET/MR scanner at UHCMC. Subjects were positioned supine on the scanner bed. The head was fixed in position using inflatable pads and an 8-channel head coil was used for data acquisition. Two fMRI scan sessions were run, one for each fraction solving strategy. They consisted of 200 echo planar imaging scans each. A high-resolution T1-weighted anatomical image and a 64-direction diffusion weighted image was also acquired. See Supplement section S1.3 for scan parameters.

### 2.7 fMRI Individual-level Analysis

FSL FEAT [36] was used to perform motion correction, spatial smoothing and apply a high pass filter to preprocess the raw data. A rigid body transform with 6 degrees of freedom was used for motion correction. Spatial smoothing was performed with a full-width-at-half-maximum Gaussian kernel of 6 mm, a high pass temporal filter of 90 s was used. All scans were then co-registered to a standard space template (MNI) and an individual level general linear model (GLM) analysis was run. The fractions were presented in a block design and a hemodynamic response double gamma function was convolved with a boxcar function to model the expected signal. We were most interested in the neural correlates of calculation so only scans during the computation time were modelled as active, not the scans during the response phase. The scans related to the response phase recorded the process of reading the possible answers and indicating their choice via a button press. The allowed response duration was kept short to ensure that calculations could not continue into this phase. Recorded response times were related to the time taken by participants to indicate their choice, not the time to do the mental calculation. Response times were not included as regressors in the GLM.

Note, one EPT subject was unable to complete the fMRI tests and a technical issue prevented one FT subject from taking the fMRI tests, hence the fMRI data reported at both individual and group level contains 13 EPT and 15 FT subjects. However, structural data (DTI and T1 images) were obtained for all 30 subjects.

### 2.8 fMRI Group Analysis

The statistical output from the individual analyses were used to perform the group level statistics using fixed effects modelling in a factorial design involving group and strategy/item type combination. Below, we focus on comparing birth group (EPT and FT) by strategy (mixed or improper), birth group by item type (Type I or Item Type II), and 3-way interaction effects. The latter consisted of within group comparisons of activations to Item Types I vs. II using the same strategy (mixed or improper), and of activations associated with the same item types (Types I and II) using different strategies. Note that strategy by item type interactions were purposely designed so that differences in difficulty levels associated with the two item types would depend on the strategy used to solve the problems. The inclusion of sex as a covariate was considered. However, given that the EPT group consisted of predominantly females (11 out of 13), the sex results were confounded with birth status and so are not reported. To reflect multiple hypotheses at least to some degree, a result was considered significant if the resulting p-value was < 0.001 which equates to a t-score > 3.1.

### 2.9 Linear SVM Classification

Linear SVM classification to further explore the activation differences between birth groups and strategy adoption was carried out. This is described in Supplement section S4.1.

### 2.10 Volumetrics and DWI analysis

MRI-based volumetrics and DWI analyses are described in the Supplement sections S5.0 and S6.0. This includes group comparisons of regional volumes and white matter connectivity.

## 3.0 Results

### 3.1 Accuracy of responses by strategy/item type

We first establish the accuracy of the performance on the fraction subtraction problems. The accuracy rates by strategy, item type (difficulty level), and strategy/item type combination are given for both the FT and EPT groups in Table 2. Note that in general accuracy is quite high, which supports the effectiveness of the training protocol, and the correct adoption of prompted strategy per item. A Wilcoxon rank sum test was used to compare the differences in accuracy rate between groups for each strategy/item type combination. Within birth groups, item types for same strategy were compared, as well as strategies for a same item type. Most comparisons were not statistically significant. Still, it was found that for both EPT and FT subjects, when using the mixed strategy, Item Type II (harder) problems had significantly lower accuracy rates compared to Item Type I (*p* = 0.007 and *p* = 0.002). Performance using different strategies on same item types were compared as well. For Item Type II, both EPT and FT had lower correctness rates (*p* = 0.049 and *p* = 0.005) for the mixed fractions strategy compared to improper fractions strategy. No significant difference was found for other item type and strategy combinations. When using the mixed strategy to solve Item Type II problems, EPT subjects had lower correctness rates compared to FT subjects (85.7 versus 71.4%, *p* = 0.050), though even the EPT had relatively high accuracy rates.

**Table 2:** Median accuracy rates for the mixed and improper fractions tasks performed during fMRI. Easy and Hard is the overall rates for the improper or mixed fractions. SD = standard deviation.

|  | **Improper**<br>**Median accuracy**<br>**rate (%) (IQR)** | **Mixed**<br>**Median accuracy**<br>**rate (%) (IQR)** |
| --- | --- | --- |
| **FT** |  |  |
| **Easy** | 100 (17.6) | 100 (14.3) |
| **Hard** | 100 (17.6) | 85.7 (17.9) |
| **EPT** |  |  |
| **Easy** | 100 (28.6) | 85.7 (0.0) |
| **Hard** | 100 (14.3) | 71.4 (25.0) |

### 3.2 Main effects of strategy and their interaction with item type

The main effect of strategy was investigated. The data were modelled as task (Item Type I and II) versus rest and both EPT and FT groups were included. The improper fraction strategy was associated with hemodynamic response in the left superior and inferior frontal gyri, paracentral lobule, precuneus and middle temporal gyrus; and right cingulate gyrus. Modelling the mixed fractions strategy in a similar way revealed hemodynamic response in left medial frontal gyrus, inferior parietal lobule and thalamus; right superior parietal lobule, superior frontal gyrus, precuneus, precentral gyrus and caudate nucleus.

The interaction of item type and strategy was also investigated. Improper fractions strategy for Item Type II (easier to solve problems) was associated with hemodynamic response in left superior and medial frontal gyri, paracentral lobule and middle temporal gyrus. Item Type I problems solved by this strategy was associated with hemodynamic response in left superior frontal gyrus and right posterior cingulate. For the mixed fractions strategy, Item Type I (easier to solve problems) showed hemodynamic response in left medial frontal gyrus, insula and fusiform gyrus; right cingulate, superior frontal gyrus, precuneus, precentral gyrus and caudate; and bilateral lingual gyrus. Hemodynamic response associated with Item Type II of mixed fractions strategy were widespread and included in left medial frontal gyrus, lingual gyrus, inferior parietal lobule, cingulate, thalamus and caudate; right superior frontal gyrus, superior parietal lobule and precuneus; bilateral precentral gyrus and posterior cingulate gyrus.

A conjunction analysis for each strategy and item types shows that much of the functional activation that differs between item types appears around the edges of the overlapping clusters. Statistical contrasts of improper strategy Item Type I versus II and Item Type II versus I were not significant. The contrast of Item Type I versus II for mixed fractions strategy showed statistical differences in left paracentral lobule, insula and precentral gyrus; right lingual and postcentral gyri. The contrast of Item Type II versus I showed greater hemodynamic response mainly in left brain areas including the precuneus, middle frontal gyrus, precentral gyrus and bilateral cingulate gyrus.

### 3.3 Between birth group differences by strategy and difficulty level

Table 3 shows between group differences in hemodynamic response for the easy and hard levels for each fraction strategy.

**Table 3:**
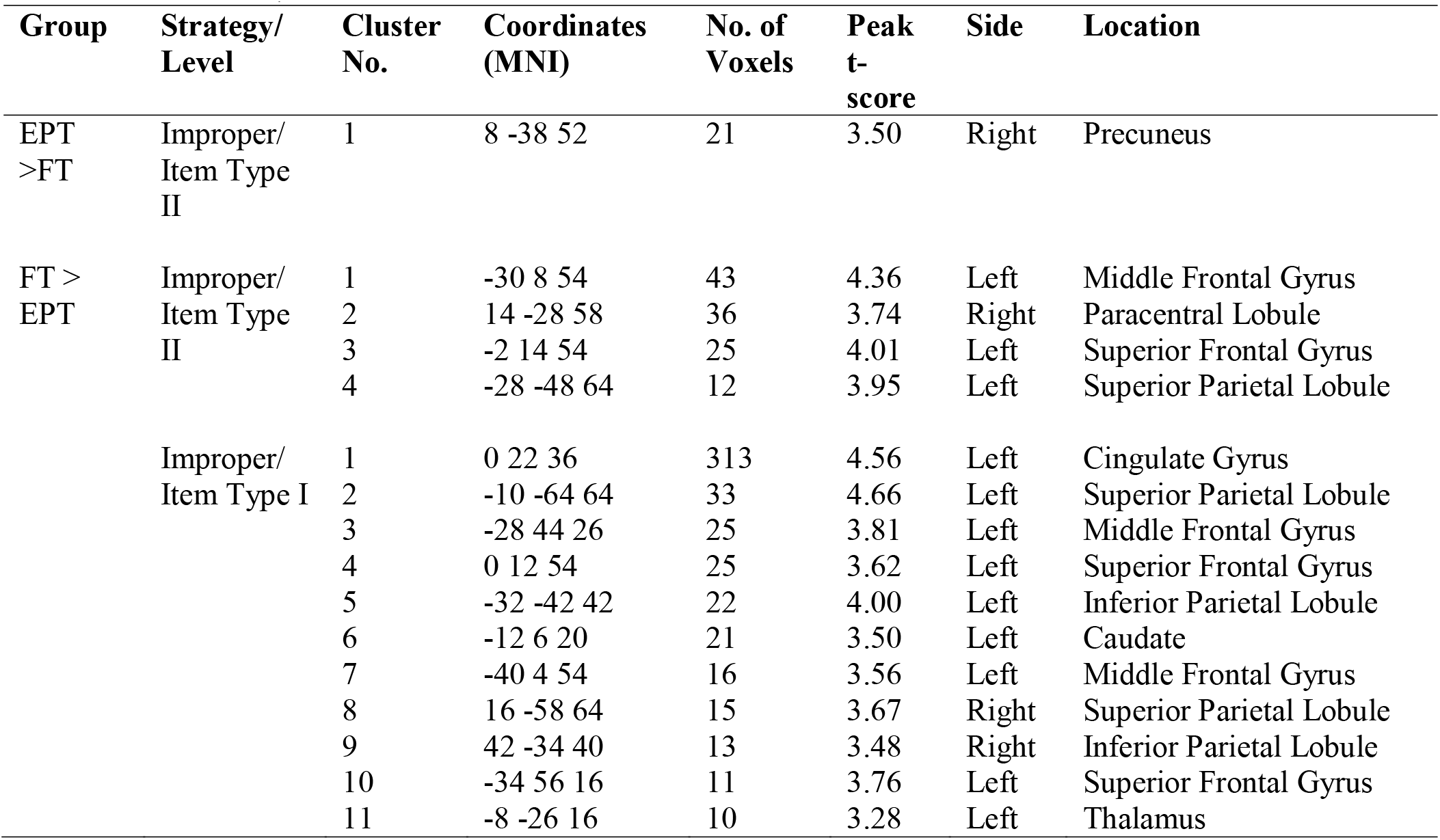
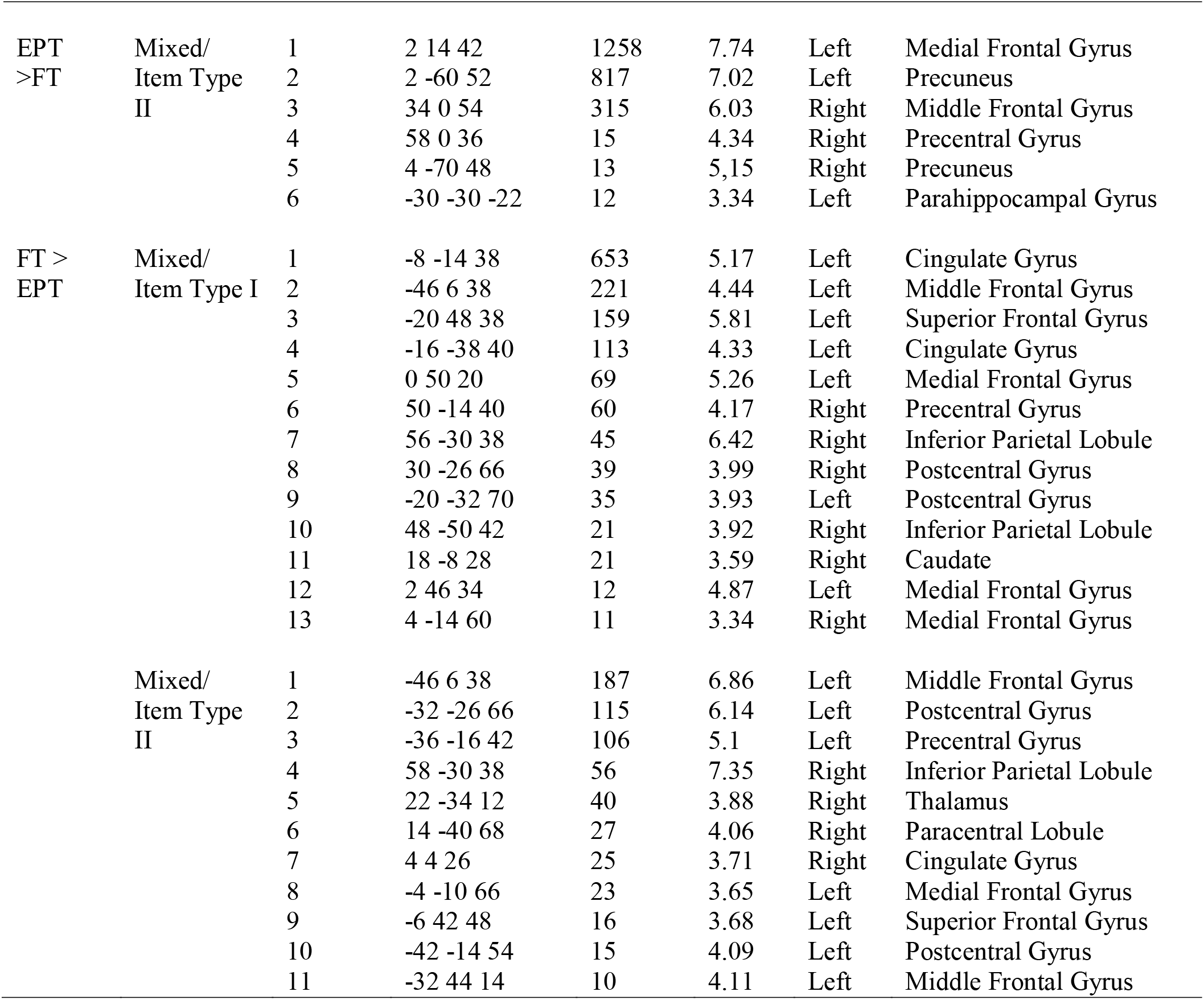
Between group differences in hemodynamic responses for Item Types for each fraction strategy. P = 0.001, minimum cluster extent = 10 voxels.

| **Group** | **Strategy/**<br>**Level** | **Cluster**<br>**No.** | **Coordinates**<br>**(MNI)** | **No. of**<br>**Voxels** | **Peak**<br>**t-**<br>**score** | **Side** | **Location** |
| --- | --- | --- | --- | --- | --- | --- | --- |
| EPT<br>>FT | Improper/<br>Item Type<br>II | 1 | 8 -38 52 | 21 | 3.50 | Right | Precuneus |
| FT ><br>EPT | Improper/<br>Item Type<br>II | 1 | -30 8 54 | 43 | 4.36 | Left | Middle Frontal Gyrus |
|  |  | 2 | 14 -28 58 | 36 | 3.74 | Right | Paracentral Lobule |
|  |  | 3 | -2 14 54 | 25 | 4.01 | Left | Superior Frontal Gyrus |
|  |  | 4 | -28 -48 64 | 12 | 3.95 | Left | Superior Parietal Lobule |
|  | Improper/<br>Item Type I | 1 | 0 22 36 | 313 | 4.56 | Left | Cingulate Gyrus |
|  |  | 2 | -10 -64 64 | 33 | 4.66 | Left | Superior Parietal Lobule |
|  |  | 3 | -28 44 26 | 25 | 3.81 | Left | Middle Frontal Gyrus |
|  |  | 4 | 0 12 54 | 25 | 3.62 | Left | Superior Frontal Gyrus |
|  |  | 5 | -32 -42 42 | 22 | 4.00 | Left | Inferior Parietal Lobule |
|  |  | 6 | -12 6 20 | 21 | 3.50 | Left | Caudate |
|  |  | 7 | -40 4 54 | 16 | 3.56 | Left | Middle Frontal Gyrus |
|  |  | 8 | 16 -58 64 | 15 | 3.67 | Right | Superior Parietal Lobule |
|  |  | 9 | 42 -34 40 | 13 | 3.48 | Right | Inferior Parietal Lobule |
|  |  | 10 | -34 56 16 | 11 | 3.76 | Left | Superior Frontal Gyrus |
|  |  | 11 | -8 -26 16 | 10 | 3.28 | Left | Thalamus |
| EPT<br>>FT | Mixed/<br>Item Type<br>II | 1 | 2 14 42 | 1258 | 7.74 | Left | Medial Frontal Gyrus |
|  |  | 2 | 2 -60 52 | 817 | 7.02 | Left | Precuneus |
|  |  | 3 | 34 0 54 | 315 | 6.03 | Right | Middle Frontal Gyrus |
|  |  | 4 | 58 0 36 | 15 | 4.34 | Right | Precentral Gyrus |
|  |  | 5 | 4 -70 48 | 13 | 5.15 | Right | Precuneus |
|  |  | 6 | -30 -30 -22 | 12 | 3.34 | Left | Parahippocampal Gyrus |
| FT ><br>EPT | Mixed/<br>Item Type I | 1 | -8 -14 38 | 653 | 5.17 | Left | Cingulate Gyrus |
|  |  | 2 | -46 6 38 | 221 | 4.44 | Left | Middle Frontal Gyrus |
|  |  | 3 | -20 48 38 | 159 | 5.81 | Left | Superior Frontal Gyrus |
|  |  | 4 | -16 -38 40 | 113 | 4.33 | Left | Cingulate Gyrus |
|  |  | 5 | 0 50 20 | 69 | 5.26 | Left | Medial Frontal Gyrus |
|  |  | 6 | 50 -14 40 | 60 | 4.17 | Right | Precentral Gyrus |
|  |  | 7 | 56 -30 38 | 45 | 6.42 | Right | Inferior Parietal Lobule |
|  |  | 8 | 30 -26 66 | 39 | 3.99 | Right | Postcentral Gyrus |
|  |  | 9 | -20 -32 70 | 35 | 3.93 | Left | Postcentral Gyrus |
|  |  | 10 | 48 -50 42 | 21 | 3.92 | Right | Inferior Parietal Lobule |
|  |  | 11 | 18 -8 28 | 21 | 3.59 | Right | Caudate |
|  |  | 12 | 2 46 34 | 12 | 4.87 | Left | Medial Frontal Gyrus |
|  |  | 13 | 4 -14 60 | 11 | 3.34 | Right | Medial Frontal Gyrus |
|  | Mixed/<br>Item Type<br>II | 1 | -46 6 38 | 187 | 6.86 | Left | Middle Frontal Gyrus |
|  |  | 2 | -32 -26 66 | 115 | 6.14 | Left | Postcentral Gyrus |
|  |  | 3 | -36 -16 42 | 106 | 5.1 | Left | Precentral Gyrus |
|  |  | 4 | 58 -30 38 | 56 | 7.35 | Right | Inferior Parietal Lobule |
|  |  | 5 | 22 -34 12 | 40 | 3.88 | Right | Thalamus |
|  |  | 6 | 14 -40 68 | 27 | 4.06 | Right | Paracentral Lobule |
|  |  | 7 | 4 4 26 | 25 | 3.71 | Right | Cingulate Gyrus |
|  |  | 8 | -4 -10 66 | 23 | 3.65 | Left | Medial Frontal Gyrus |
|  |  | 9 | -6 42 48 | 16 | 3.68 | Left | Superior Frontal Gyrus |
|  |  | 10 | -42 -14 54 | 15 | 4.09 | Left | Postcentral Gyrus |
|  |  | 11 | -32 44 14 | 10 | 4.11 | Left | Middle Frontal Gyrus |

**Table 4.**
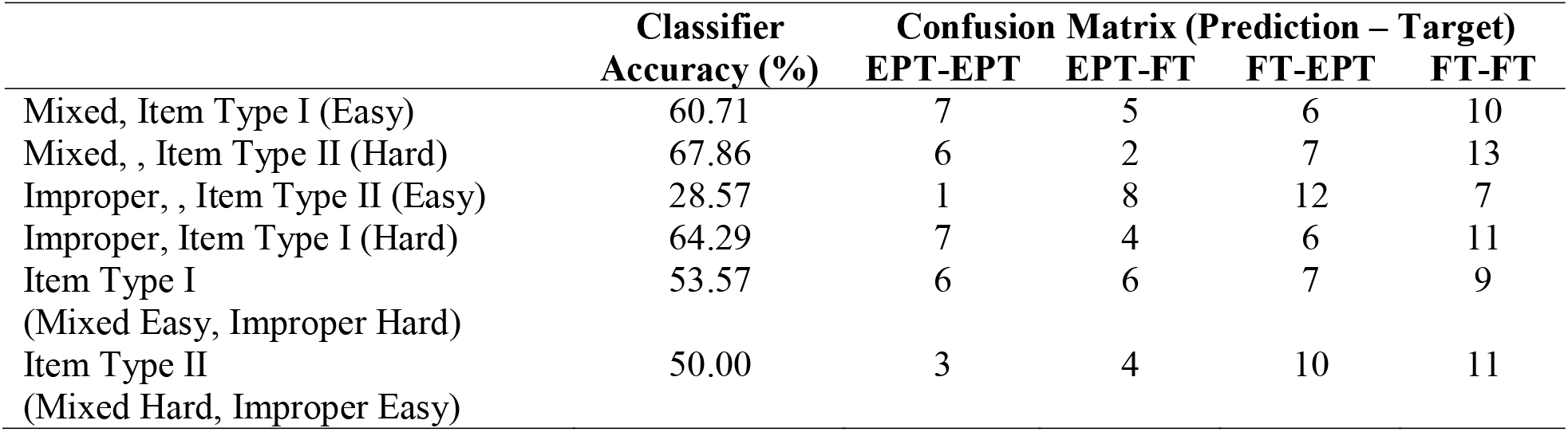
Accuracies and confusion matrices from SVM classification of birth status.

#### 3.3.1 Improper strategy

Comparison of the EPT vs FT groups on the less difficult items (Item Type II) for the improper strategy vs rest revealed greater hemodynamic response in right precuneus in the EPT group compared to the FT group (contrast EPT vs FT). The FT group showed greater hemodynamic response in the left middle and superior frontal gyri, superior parietal lobule and right paracentral lobule (contrast FT vs EPT). For Item Type I (hard) vs rest, we observed no areas displaying greater hemodynamic response in the EPT group compared to the FT group. For the more difficult Item Type I, the FT group showed greater hemodynamic response in left cingulate gyrus, middle and superior frontal gyri, caudate and thalamus, and bilateral superior and inferior parietal lobules compared to the EPT group. Locations for each contrast are shown in Figure 3 and listed in Table 3. A conjunction analysis of EPT and FT group activity is shown in Figure 4. These results are suggestive of the EPT group demonstrating either widespread under-activation during the task compared to FT subjects or it may be due to more variability in brain responses across subjects. Closer inspection of the variance between the groups showed that the EPT group variance is almost twice as much as the FT group – for example FT improper Item Type I: mean voxel level variance = 222.6 (SD 151.6) and EPT improper Item Type I: mean voxel level variance = 428.2 (SD 327.2). Across all strategies and item types, the variance in the EPT group remains consistently around twice the variance in the FT group.

**Figure 3:**
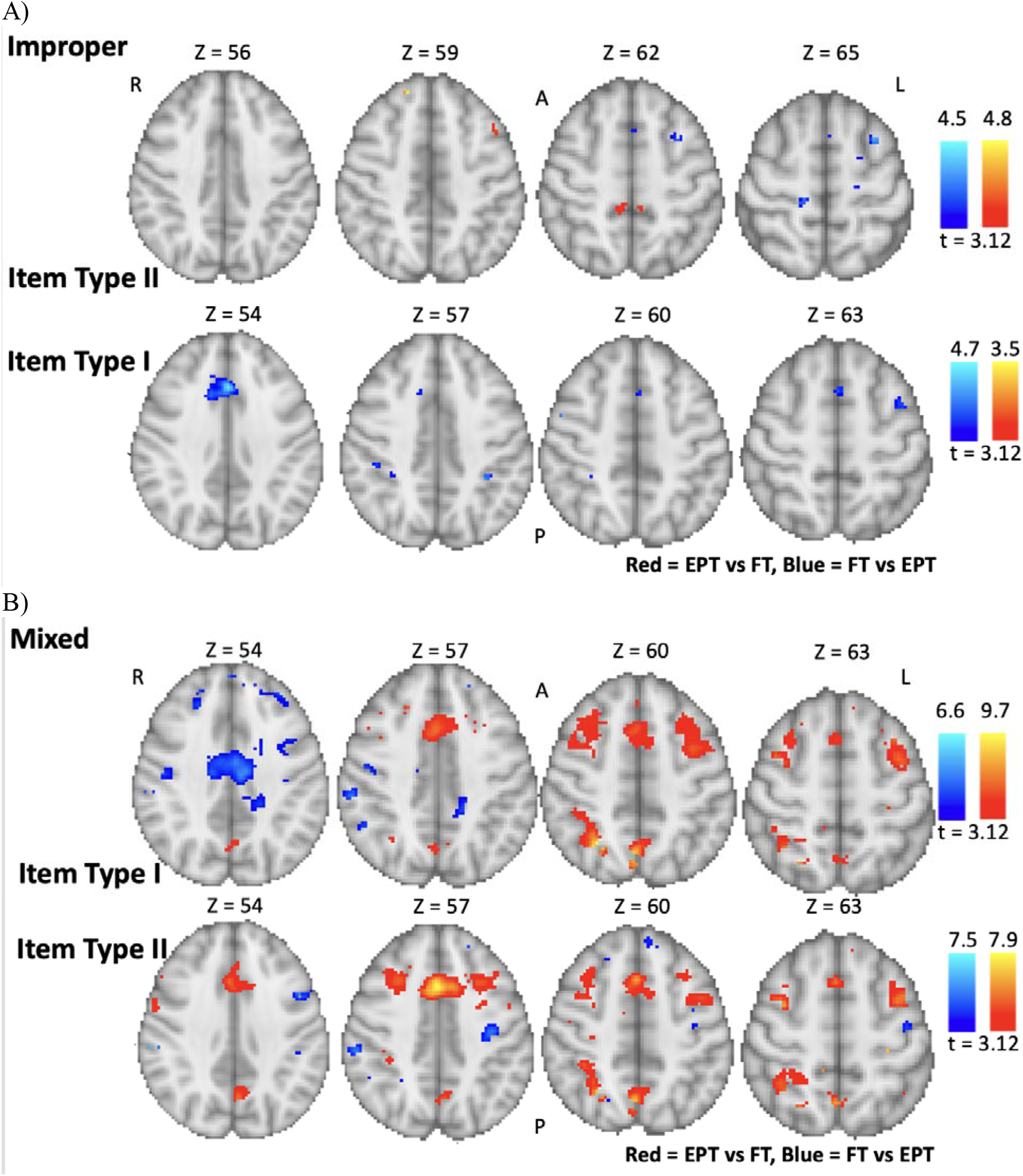
Hemodynamic responses associated with improper and mixed fractions for both groups. Contrasts are easy EPT vs FT, easy FT vs EPT, hard EPT vs FT, and hard FT vs EPT. Improper fractions (A) and mixed fractions (B) locations are shown separately. L = Left, R = Right, A = Anterior, P = Posterior. Regions include - left side: precuneus, superior, medial and middle frontal gyri, superior and inferior parietal lobule, pre- and post-central gyri, insula, cingulate gyrus, parahippocampal gyrus, caudate nucleus and thalamus. Right side: paracentral lobule, precuneus, pre- and post-central gyri, inferior and superior parietal lobule, superior, medial and middle frontal gyri, cingulate, caudate nucleus and thalamus.

**Figure 4:**
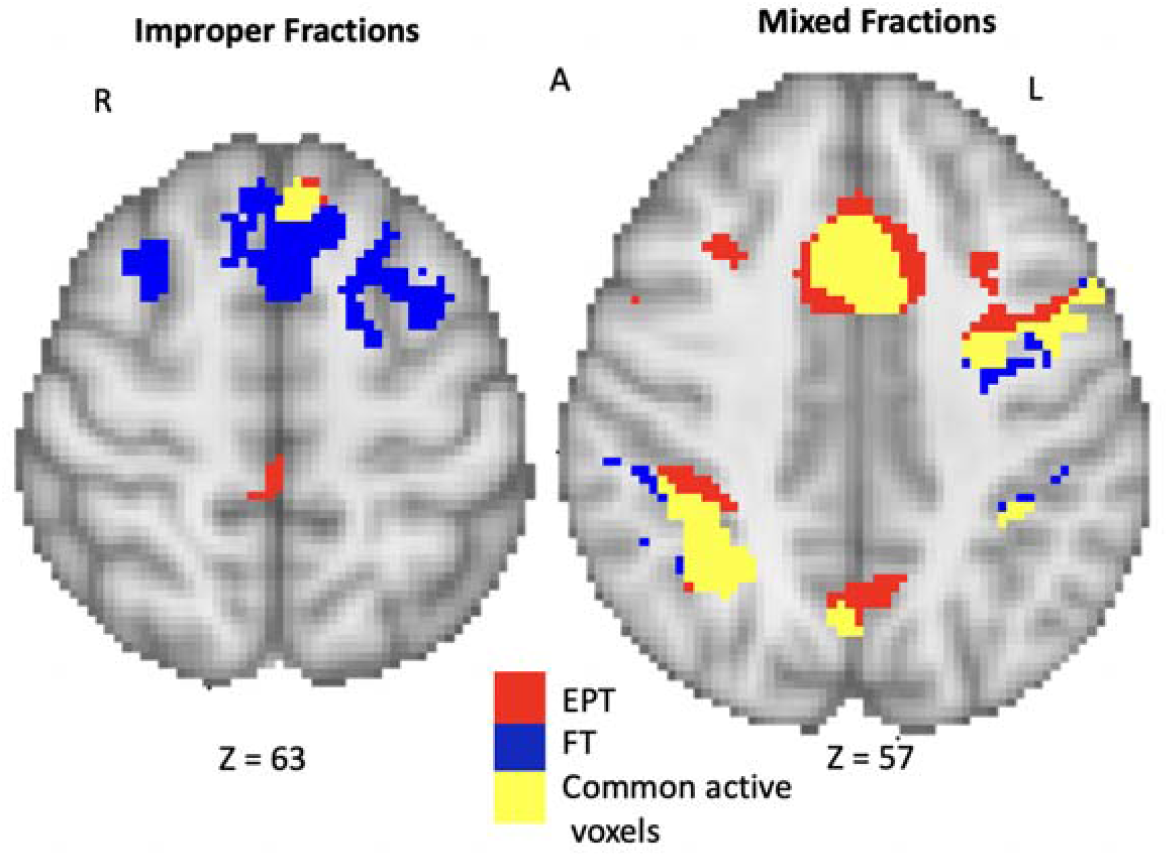
Conjunction analysis of EPT and FT group results for improper and mixed fractions. The EPT (red) and FT (blue) results have been overlaid. Shared voxels between EPT and FT are shown in yellow. Improper fractions and mixed fractions include both Item types I and II. The brain activity is overlaid on the MNI template brain. L = left, R = right, A = anterior.

#### 3.3.2 Mixed strategy

In group comparisons of the less difficult items (Item Type I) versus rest for the mixed strategy data, the EPT group compared to the FT group showed greater hemodynamic response in left insula, middle and medial frontal gyri; right superior frontal gyrus, cingulate, superior parietal lobule and precuneus. In contrast, the FT group compared to the EPT group showed greater hemodynamic response in left cingulate, middle and superior frontal gyri; right precentral gyrus, inferior parietal lobule, caudate nucleus; bilateral medial frontal and postcentral gyri. For Item Type II (hard) versus rest, the EPT group compared to the FT group showed greater hemodynamic response in left medial frontal gyrus and parahippocampal gyrus; right middle frontal and precentral gyri; bilateral precuneus. For the easier Item Type II, the FT group had greater hemodynamic response in left middle, medial and superior frontal gyri, pre- and post-central gyri; right inferior parietal lobule, cingulate, paracentral lobule and thalamus compared to the EPT group. The conjunction analysis for mixed fractions Item Types I and II in Figure 5 shows that much of the activity that is different between the groups appears around the edges of the overlapping clusters where statistical certainty is the lowest. These areas coincide with those listed above.

**Figure 5:**
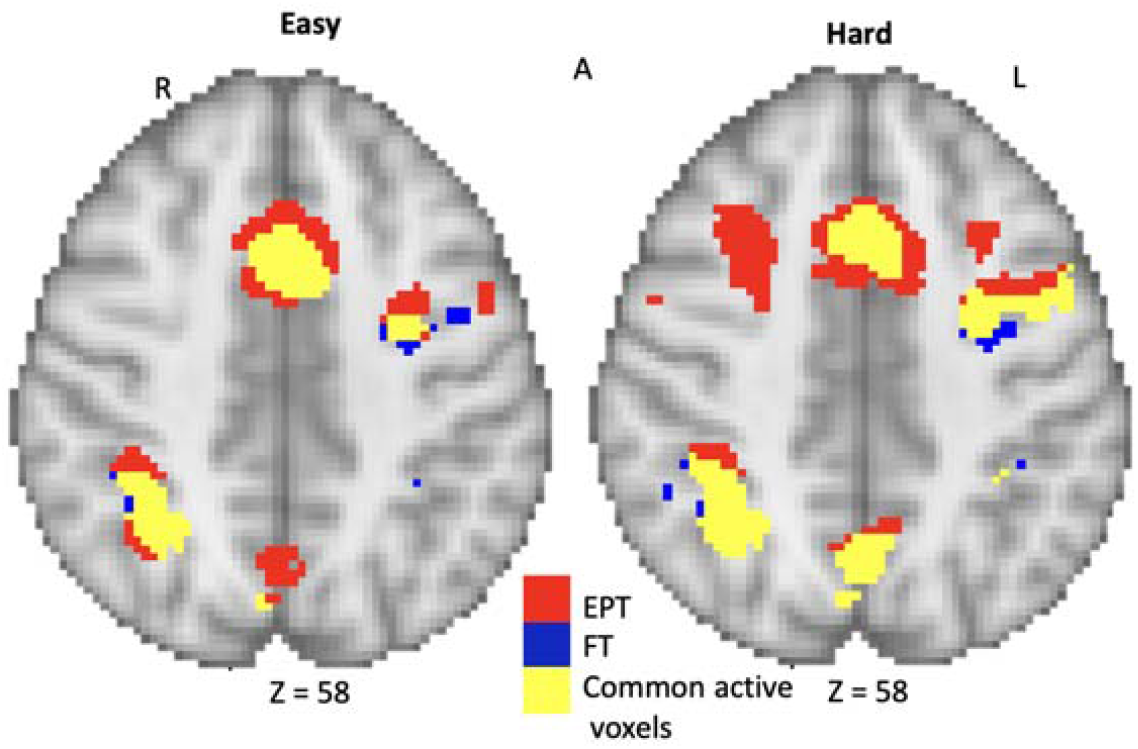
Conjunction analysis of Item Types I and II for FT and EPT for mixed fractions. The EPT (red) and FT (blue) results have been overlaid for each Item Type. Shared voxels are shown in yellow. The brain activity is overlaid on the MNI template brain. L = left, R = right, A = anterior.

**Figure 6:**
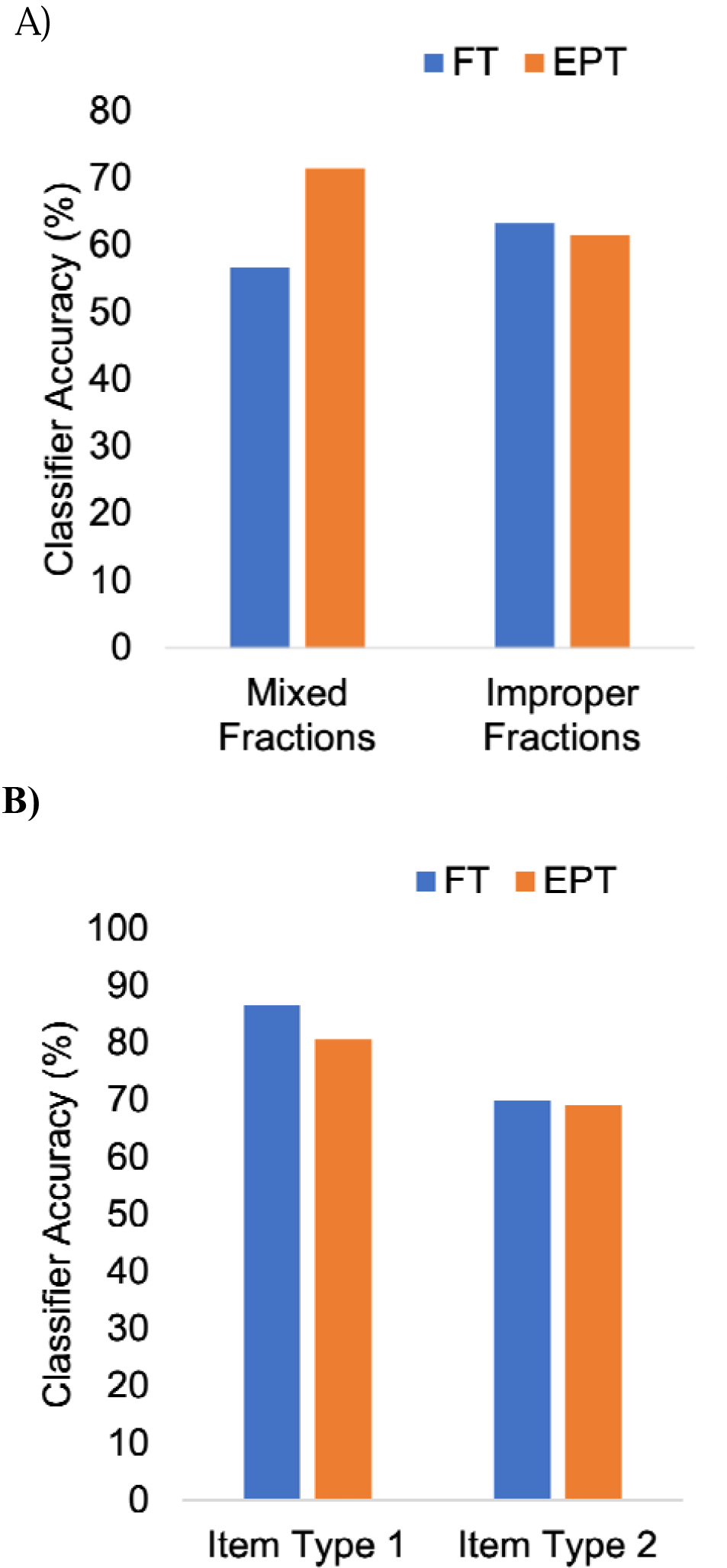
Accuracies of group-level SVM classification of difficulty level (A) and strategy (B) from both FT and EPT groups. A leave-one-out cross validation approach was used for all SVM analyses.

## 4.0 Discussion

This study explored differences in hemodynamic response between extremely preterm and FT born teens associated with using two strategies to solve fractions, where cognitive loads varied by item type and strategy. A main hypothesis of the study was that among the specific strategy and item type combinations, activation differences between birth groups would be particularly pronounced for the harder versus easier difficulty levels of a given strategy.

Overall, the estimated main effects of strategy indicated hemodynamic responses across widespread areas of the brain associated with mathematical processing. Mixed fractions strategy was associated with the highest number of active voxels and with a pattern of functional activation that was distinct from the improper fractions strategy. Mixed fractions related hemodynamic responses were located in fronto-parietal areas, while improper fractions hemodynamic responses were centered around the precuneus and temporal gyrus. The hemodynamic response of the mixed fractions strategy is consistent with that reported in the literature for mathematics tasks [8, 9].

In terms of group differences, the interaction between strategy and item type showed areas of greater hemodynamic response specific to each group for each condition, with the exception of Item Type I improper fractions strategy, where the EPT did not show greater functional activation compared to the FT group. The areas of increased hemodynamic response for all other conditions were widespread in fronto-parietal areas and along the midline and temporal gyri and by birth group, strategy and item type. Efforts to solve both item types (I and II) using the mixed fractions produced similar patterns of activity for both groups (see Supplement section S2.0, within group analyses). In comparison, functional activations for this strategy in the EPT group compared to the FT group were greater in bilateral insulae and middle frontal gyri. The different networks for each strategy may be due to the different processes involved with the calculations (see Figure 1). The mixed fraction strategy can have a greater cognitive load attributable to a higher demand for number memorization necessary to achieve the answer (i.e., the separate answers for the whole number and fractions must both be remembered and one must be kept in memory while the other is calculated), particularly for Type II items (harder level). This finding may also reflect a relatively higher level of demand on executive function. In contrast, when using the improper fractions strategy, each calculation at a given step is used in the next step until the final result is obtained. Some degree of working memory is thus needed to apply the improper fractions strategy, particularly for Type I items (harder level). However, this approach may not require the same magnitude of retention and manipulation as the mixed fractions strategy. The difference between the strategies may also be due to more familiarity with the improper fractions strategy which is often taught in schools. In support of this possibility, Tenison and colleagues found that unfamiliar mathematics problems, compared to more familiar ones, took longer to solve and were accompanied by increased brain activity in the intraparietal sulcus [37].

Group differences in patterns of functional activation may be associated with several factors. The EPT subjects generally had lower working memory function (see WRAML2 scores in Table 1) and they may have adopted different cognitive strategies to compensate for this weakness, resulting in different areas of neural recruitment. An alternative explanation is that there may be alterations in brain structure. Consistent with the group differences in brain structure observed in this study (discussed below), past research has revealed alterations in grey and white matter volumes and altered folding patterns of the cortical gyri in individuals born EPT [38, 39]. The pattern of functional activity observed here may also be reflective of compensation for reduced capacity or neural efficiency in the areas typically used by FT control subjects. A previous study found evidence for neural compensation in adults with preterm birth on a visual learning task [40]. Further evidence in support of compensatory processes, as manifest in “cognitive reserve,” is provided by findings indicating age differences between younger and older adults in the level of task difficulty at which patterns of neural activation change from those observed while doing less difficult problems [41]. Greater variability in brain response at the individual level may also have contributed to activation differences between the EPT and FT groups. Greater variability in functional brain activity has been found among those with developmental dyscalculia [42] and those born EPT [43]. The source of the greater variability may arise from differences in the brain structure associated with preterm birth [38, 39], but may also reflect different developmental trajectories related to atypical patterns of maturation and neuronal pruning [44, 45]. One study found evidence of accelerated brain maturation and earlier neuronal pruning in individuals with EPT compared to those born FT [46].

The SVM-based classifications with the highest accuracies were observed for strategy classification based on item type (i.e., difficulty level). This higher classification accuracy (> 80%) is seen for both groups. These findings suggest BOLD signals that differentiate strategy may be shared across groups and are consistent with the group-level GLM activation pattern results, which indicate different neural activation patterns by strategy. Classification rates for distinguishing difficulty levels were higher for the EPT group than for the FT participants, particularly for the mixed fractions strategy (71.43% versus 56.67%). This finding provides further evidence that the EPT participants may be straining to solve the more difficult Type II items with this strategy. These results are consistent with other findings from this study, including group differences in accuracy, GLM-based activation patterns, and DWI-based connectivity, indicating that group differences were most pronounced when participants were solving more difficult problems.

The above-noted group differences in patterns of brain functional activations and structure suggest that some youth with EPT birth may engage different brain networks than FT youth in solving fraction problems. These findings suggest a need to tailor instructions in fraction problem-solving, and perhaps other aspects of mathematics, to individual differences. In solving problems relying on brain networks similar to those engaged by FT youth, similar approaches may be justified. However, the present findings indicate that at least some EPT adolescents may engage different brain networks, depending on the strategy employed in solving a problem, the type of item, and the cognitive competencies of the learner. One implication of these findings for mathematics instruction is the support they provide for adapting problem-solving strategies in ways that are biologically informed. Although procedures for making these modifications are to be determined, the present results suggest the need for the student to recognize both their individual cognitive competencies and the characteristics of the mathematics problems, such as their cognitive load and the corresponding respective efficiencies of problem-solving strategies. Alternatively, efforts could be made to strengthen some of the processes that contribute to problem-solving. An example of these efforts as applies to fraction problem-solving would be to increase the automaticity of retrieval of arithmetic facts as a means for compensating for weaknesses in working memory. In the Supplement section S3.0, it was found that several areas were more likely to be functionally active in the EPT group than the FT group, including the left parahippocampal and bilateral medial frontal gyri, as well as a lack of functional activation in the precuneus. The precuneus has been studied as a functional hub between multiple interacting brain networks and supports visuospatial imagery and attention activation [47], skills necessary for mathematical calculations.

We also compared the EPT and FT groups on measures of brain structure using volumetrics and DTI analysis (see Supplement sections S5.0 and S6.0). Volumetric analysis revealed reduced volume in the left lingual and parahippocampal gyri in the EPT compared to the FT group. DTI analysis revealed several altered connections for these youth, particularly involving the left parahippocampal gyrus, left visual area and right superior parietal lobule. The findings indicating alterations in the EPT group in brain volumes and connectivity are in line with other studies in very preterm individuals (GA < 33 weeks) [38, 39, 48, 49]. Increased volumes and abnormal patterns of structural connectivity may reflect alterations in brain growth that occur in response to, or in compensation for, the above-noted decreases in brain volumes. For example, stronger connectivity between some brain regions may compensate for weaker connectivity between other regions. The overlap we found between the volumetric and DTI analyses suggests structural changes in the left parahippocampal gyrus and right superior parietal lobule. Both regions had reduced white matter volume and structural connectivity. The superior parietal lobule is involved with action processes and visuomotor functions, attention, reasoning, spatial perception, visual perception and working memory [50]. The function of the parahippocampal gyrus has been linked to retrieval fluency in 7- to 9-year-old children [51]. Reduced grey and white matter volume in this area may impact fluency in retrieval of math facts and suggests that EPT individuals may need to invest more effort in making mental computations. The volumetric analysis also suggested that the EPT group had reduced grey and white matter volume in the left lingual gyrus. The left lingual gyrus has been linked to abilities in visual memory [52], a component of the working memory system [53]. Reduced volume in both the grey and white matter of the lingual gyrus areas raises the possibility of deficits in aspects of the working memory system supported by that region.

Major study limitations are the small sample size and questions regarding sample representativeness. Participants were recruited from a larger sample of youth assessed several years prior to this study. Many of the families could not be located. Although the EPT and FT groups did not differ in age, most youth in the EPT group were female; thus, the sample was not representative of the broader regional or national population of EPT youth. Further MRI studies with larger and more representative samples are needed to confirm our results, and to continue to explore how heterogeneity in cognitive outcomes of EPT birth may be associated with variability in neural activations that we observed at the group level.

## 5.0 Conclusion

The effects of being born EPT on the brain are multifactorial and complex. The present study explored brain function and structure in those born EPT through the lens of mathematical strategies used in performing mental computation of fraction problems. We found both group and strategy-dependent differences in neural activations to fraction problem-solving. The findings offer support for considering individual differences among EPT in neural structure and function, along with the specific demand characteristics or mathematics problems, in developing more targeted approaches to mathematics instruction. This information may inform pedagogical practices and allow educators to tailor mathematical approaches adapted to the cognitive and neural profiles of EPT individuals. Particularly, hemodynamic response patterns associated with different problem-solving strategies in youth born EPT compared to FT youth could serve as potential biomarkers of approaches to mathematics instruction that may be most appropriate for youth born EPT. The present findings suggest consideration of instructional approaches that focus on strategies with relatively lower cognitive loads and on emphasizing flexibility in strategy adoption.

## Declaration of conflicts of interest

All authors declare no conflicts of interest.

SC – Study design, analysis and interpretation of data, drafting of manuscript

MB – Study design, analysis and interpretation of data, review of manuscript

AI – Study design, analysis and interpretation of data, review of manuscript

BM-D – Study design, analysis and interpretation of data, review of manuscript

GW – analysis and interpretation of data, review of manuscript

WC – Study design, analysis and interpretation of data, software development, review of manuscript

JF – Study design and software development, review of manuscript

JZ – Study design, analysis and interpretation of data, review of manuscript

MW – analysis and interpretation of data, review of manuscript

SYY – analysis and interpretation of data, review of manuscript

AB – Study design, data collection, review of manuscript

ER – Study design, analysis and interpretation of data, review of manuscript

CG – Study design, analysis and interpretation of data, review of manuscript

NM – Study design, analysis and interpretation of data, review of manuscript

LT – Study design, analysis and interpretation of data, review of manuscript

WHK - Study design, analysis and interpretation of data, review of manuscript

YS – study design, interpretation of data, review of manuscript

CN – interpretation of data, review of manuscript

HGT – Study design, analysis and interpretation of data, drafting of manuscript

CT – Study design, analysis and interpretation of data, drafting of manuscript

## Funding

This study was supported by Phillips Healthcare and the National Science Foundation (Award number: 1561716).

## Acknowledgements

None

## Data availability

The raw data supporting the conclusions of this article will be made available by the authors, without undue reservation.

## Notes

### Competing Interest Statement

The authors have declared no competing interest.

### Summary of Updates

General editing of the text to improve the flow for the reader and make the document more succinct. Information from some sections has been moved to the Supplement document - 2.4, 2.5, 2.6, 2.9 and 3.4.

