## supplementary material for "Fractions strategy differences in those born extremely preterm"

**S1.0 Background info:**

S1.1 Fraction Strategy Training

The multi-step training program began with review of basic concepts followed by (1) instruction and practice in converting whole and mixed numbers to proper and improper fractions, subtraction of simple proper fractions, and written subtraction problems, using each of the two strategies in order of increasing difficulty; 2) practice and review in mental problem solving, using two strategies (the “improper fraction” and “mixed fraction” approach) for solving problems that are easier or harder depending on whether the items are Type I or Type II ; and 3) computer administration of similar sets of problems to prepare participants to solve fraction problems in the same format that was used for subsequent in-scanner presentations. During training with written problems, the examiner taught each of the two problem solving strategies explicitly and had the subject verbalize the strategy after solving each problem. Practice in doing the problems mentally also included the requirement that subjects verbalize the strategy used after giving their answers. For the computer administrations, subjects were first presented with a problem on the screen and asked to provide the answer verbally as soon as they could while pressing the space bar to indicate their time to solve the problem. They were then asked to verbalize the strategy used to solve that problem. Problems were limited to those with common denominators and subtraction of relatively simple fractions to ensure that subjects could be taught the problem-solving strategies to a high level of accuracy and that in-scanner mental calculations would be feasible.

S1.2 fMRI Stimulus protocols

The visual stimulus was presented using an in-house custom written program that was developed using Python (Python Software Foundation, <https://www.python.org/>) and libraries from PsychoPy - an open source visual presentation program [1-3]. The Cedrus Lumina controller was used to integrate all the signals. The program connected to the controller box which received the subject responses as well as a trigger pulse from the MRI scanner. The trigger pulse was outputted every scan (every 3 seconds) and was used to synchronize the stimulus to the scan acquisition. On the experimenter’s computer screen was displayed a Supervisor Window that tracked the current block number being presented, how many remaining blocks there were and when the subject responded. The software has been made available from the Bitbucket repository: <https://bitbucket.org//tatsuoka-lab/fmri-presentation>.

S1.3 MR Imaging Parameters

Echo planar imaging scans were acquired with the following parameters: TR = 3.0 s, TE = 35 ms, voxel resolution = 1.797 x 1.797 x 4 mm (matrix 128 x 128), 36 slices in total provided full brain coverage, and flip angle was 90°. A SENSE P reduction factor of 2 was implemented and scans were acquired in an ascending interleaved fashion. Two fMRI scan sessions were run, one for each fraction solving strategy. They consisted of 200 scans each. A high-resolution T1-weighted anatomical image was also acquired using a magnetic preparation gradient-echo sequence (3D IR TFE) with the parameters: TR = 7.5 ms, TE = 3.7 ms, voxel resolution = 1 x 1 x 1 mm, number of slices = 200 slices and flip angle = 8°. A 64-direction diffusion weighted image with a FOV of 224 x 224 producing a final in-plane resolution of 1.75 x 1.75 mm^2^ and 2 mm slices. A TR of 7415 ms was used. Full-brain coverage was achieved with 60 slices. One B_0_ image was acquired, and the b factor was 1000.

S2.0 fMRI Results

S2.1 Within birth group generic brain activation maps

We investigated the contrasts related to differences in strategy for each item type, and differences in item types for each strategy. The latter contrasts assess how hemodynamic responses differ across difficulty levels, as in notions of cognitive reserve [4-6].

S2.1 Mixed and improper strategies.

No spatial overlap was evident in hemodynamic responses for the mixed compared to improper strategies for the EPT group. Activations for these two strategies slightly overlapped for the FT group, with 146 active voxels in total divided across 3 clusters located in bilateral superior and middle frontal gyri. Both the FT and EPT groups demonstrated similar networks for each problem-solving strategy, see Figure S1. The mixed fraction network included the intraparietal sulcus and medial and middle frontal gyri. The neural correlates of the improper fractions were located around the precuneus/paracentral lobule and frontal pole area. Generally, fewer active voxels were detected for the improper method across item types and the maximum t-scores were lower than those for the mixed method – improper fractions maximum t-scores remained below 5.0 across all conditions; mixed fractions maximum t-score ranged from 5 – 14 across all conditions. More detailed exploration of the data is given in the following sections.

S2.2. Improper strategy at two levels of difficulty.

The location of the generic brain activations for the two difficulty levels (easy and hard) for improper and mixed fractions strategies among EPT and FT participants are listed in Table S1. Table S1 also displays within group differences in activity between easy and hard difficulty levels by fraction strategy. For improper fractions easy condition, the EPT group activity local maxima were in the superior frontal gyrus, precuneus and parahippocampal gyrus. The FT group activity local maxima were in middle and superior frontal gyri, paracentral lobule and postcentral gyrus. For improper fractions, the contrasts of Item Type I and II (Easy vs Hard and Hard vs Easy) for the EPT group showed no significant differences. For the FT group, a contrast of Item Types II vs I (Easy vs Hard) showed more activity during the easy task in left medial frontal, precentral and postcentral gyri. The contrast Hard vs Easy showed no significant differences for the FT group.

S2.3 Mixed strategy at two levels of difficulty.

The hemodynamic responses associated with the mixed fractions strategy differed from those associated with improper fractions; they were more frontoparietal, and activity was more widespread by comparison to improper fractions (Figure S1). The activity for Item Types I (easy) and II (hard) are listed in Table S1 and presented in Figure S2 for both groups. In Figure S3, we see substantial overlap of active areas between easier and harder levels for both the FT and EPT groups: 49% of voxels for FT and 61% for EPT. For the EPT group, the contrast of Item Type I vs II for the strategy showed that the insula, transverse temporal gyrus, inferior parietal lobule, cingulate gyrus, lingual gyrus, precentral gyrus and paracentral lobule were more active during Item Type I. Locations more active during Item Type II were cingulate gyrus, precuneus, precentral and middle frontal gyri. The contrast of Item Type I vs II for the FT group showed the thalamus, inferior parietal lobule, cingulate gyrus, parahippocampal and pre- and post-central gyri areas were more active during the easier level. Locations more active for harder items compared to easier ones (Hard vs Easy) for this strategy were frontal areas and the precuneus.

S2.4 Item Type I problems solved using the mixed vs. improper strategy.

For the FT group, Item Type I problems (harder level for improper fractions vs easier level for mixed fractions strategies) showed increased activations associated with improper fractions in left posterior cingulate and middle frontal gyri, and right paracentral lobule and superior and middle frontal gyri. The contrast of mixed easy vs improper hard showed activity associated with the mixed fractions strategy in left cingulate and precentral gyri, and right superior parietal lobule, precuneus and posterior cingulate gyrus. In the EPT group, Item Type I problems (improper hard vs mixed easy) revealed a more widespread pattern of activity associated with the improper fractions. These included left paracentral and superior parietal lobules; and bilateral superior frontal gyrus and posterior cingulate gyrus. For mixed easy vs improper hard, activity was in left medial frontal gyrus, superior temporal gyrus and superior frontal gyrus; right superior parietal lobule, precuneus, middle frontal gyrus; and bilateral insula and precentral gyrus.

S2.5 Item Type II problems solved using mixed vs. improper strategies.

For Item Type II comparisons (easier level for improper fractions vs harder level for mixed fractions strategies), the EPT group showed activity related to improper fractions in precuneus, superior, middle and inferior frontal gyri, posterior cingulate and postcentral gyri; right inferior parietal lobule; and bilateral precentral gyri and parahippocampal gyri. The same contrasts in the FT group revealed increased activity associated with the improper fractions in left superior and middle frontal gyri. Activity associated with mixed fractions in the EPT group was located in left medial frontal gyrus and inferior parietal lobule; right lingual gyrus and insula; bilateral superior parietal lobule, thalamus, superior and middle frontal gyri and precentral gyrus. In the FT group, activity associated with the mixed fractions was in left precentral gyrus, middle frontal gyrus and insula; right inferior parietal lobule and precuneus; and bilateral cingulate and lingual gyri.

**Figure S1:** Hemodynamic responses associated with the improper and mixed fraction solving strategies. Item Types I and II (Easy and hard) were modelled together as one variable. Results for the full term (FT) group are shown on the left, for the extremely preterm (EPT) group on the right. For both groups, improper fractions are shown in red and mixed fractions are shown in blue. The brain activity is overlaid on the MNI template brain, slice z = 64 is shown, p = 0.001. L = left, R = right, A = anterior and P = posterior.

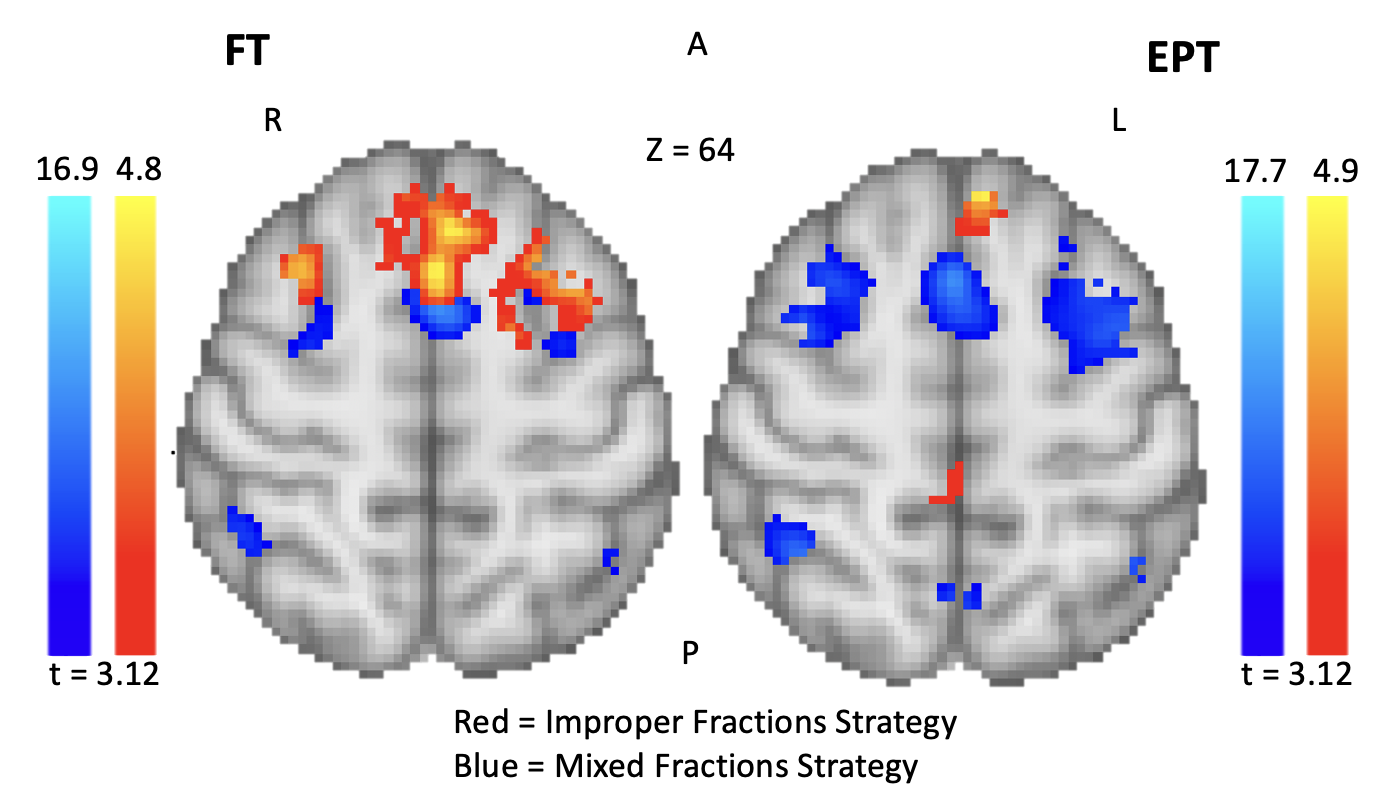

**Figure S2:** Hemodynamic responses associated with mixed fractions for the contrasts Item Type I and II (Easy vs Hard and Hard vs Easy). FT (top row) and EPT (bottom row). L = Left, R = Right, A = Anterior, P = Posterior. Regions include - left side: cingulate, inferior parietal lobule, middle frontal gyrus, postcentral gyrus, insula, paracentral lobule, precuneus and precentral gyrus. Right side: cingulate, superior frontal gyrus, precuneus, precentral gyrus, parahippocampal gyrus, thalamus, lingual gyrus and transverse temporal gyrus.

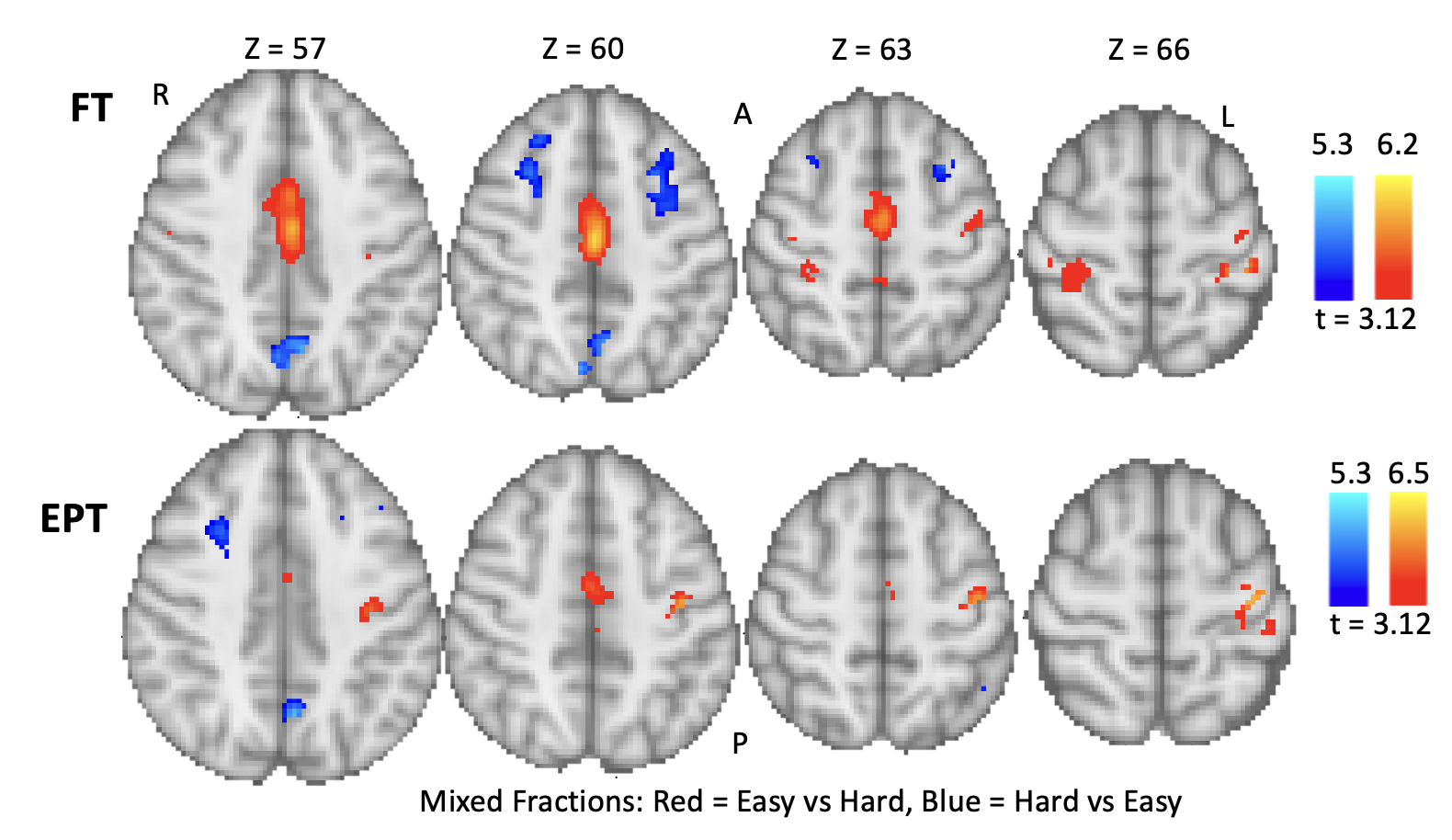

**Figure S3:** Conjunction analysis of Item Types I and II for improper and mixed fractions. Easy level (red) and hard levels (blue) have been overlaid and those voxels which are in common between the difficulty levels are shown in yellow. Top row - FT group results, bottom row – EPT group results. The brain activity is overlaid on the MNI template brain. L = left, R = right, A = anterior and P = posterior.

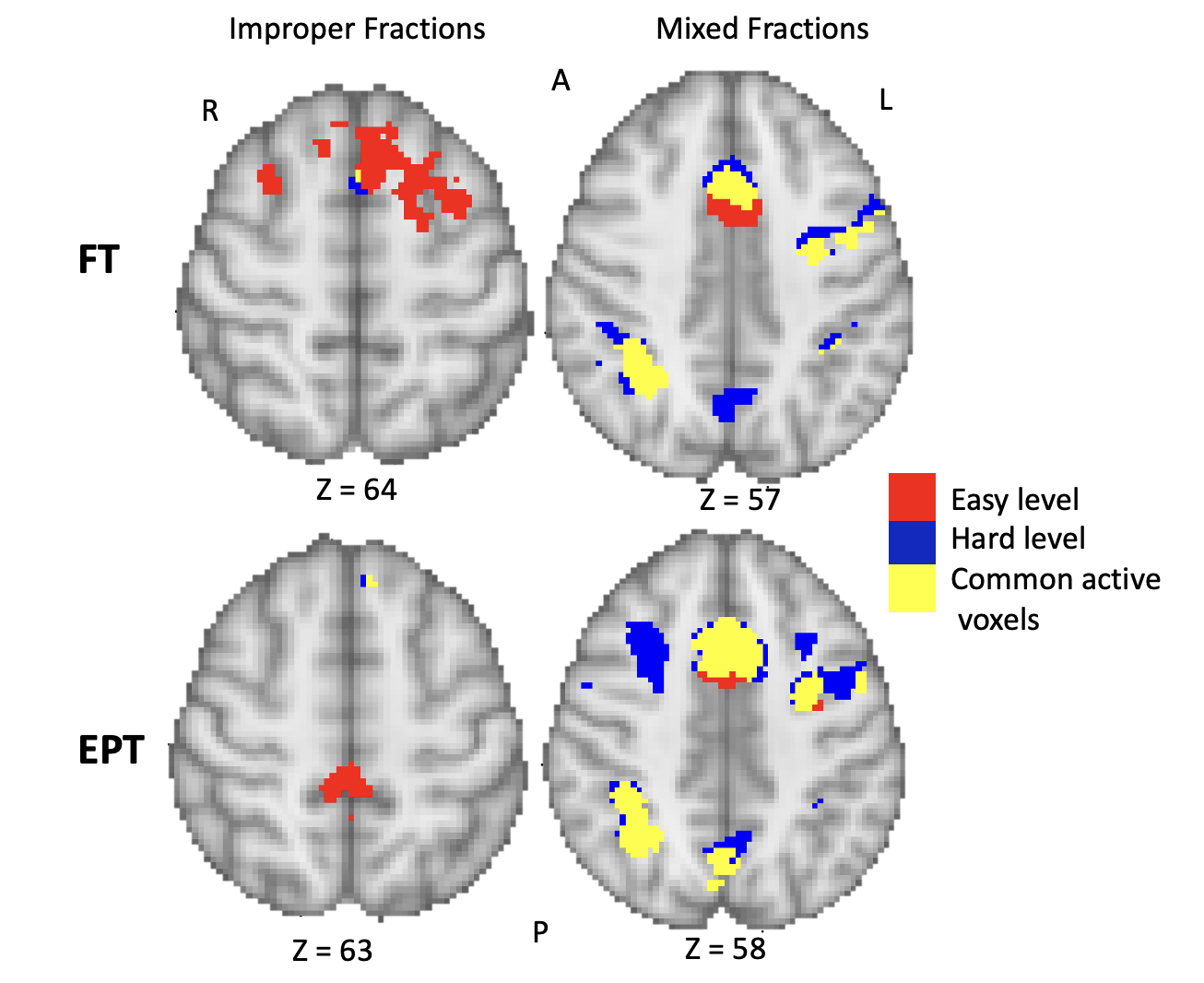

**Table S1:** Local maxima of hemodynamic responses for the EPT and FT groups during the improper and mixed fractions strategy. The results are broken down by Item Type. P = 0.001, minimum cluster extent = 10 voxels.

| **Group** | **Strategy/Level** | **Cluster No.** | **Coordinates (MNI)** | **No. of Voxels** | **Peak t-score** | **Side** | **Location** |
| --- | --- | --- | --- | --- | --- | --- | --- |
| EPT | Improper/  Item Type II (Easy) | 1  2  3 | 0 -32 52  -24 -44 -12  -4 44 46 | 129  106  13 | 4.08  3.95  3.45 | Left  Left  Left | Precuneus  Parahippocampal Gyrus  Superior Frontal Gyrus |
|  | Improper/  Item Type I (Hard) | 1  2  3 | -4 40 50  -6 32 56  2 -26 66 | 14  12  11 | 3.79  3.65  3.35 | Left  Left  Left | Superior Frontal Gyrus  Superior Frontal Gyrus  Paracentral Lobule |
| FT | Improper/  Item Type II | 1  2  3  4  5 | -30 10 58  -4 -30 62  12 -28 64  12 24 56  -22 -20 72 | 466  118  101  20  13 | 4.60  3.72  3.82  3.42  3.55 | Left  Left  Right  Right  Left | Middle Frontal Gyrus  Paracentral Lobule  Paracentral Lobule  Superior Frontal Gyrus  Postcentral Gyrus |
|  | Improper/  Item Type I | 1  2  3  4 | 2 12 62  52 0 48  -34 52 18  -48 8 18 | 53  40  39  15 | 4.16  4.52  4.37  3.61 | Left Right  Left  Left | Medial Frontal Gyrus  Precentral Gyrus  Middle Frontal Gyrus  Inferior Frontal Gyrus |
| FT | Improper/  Item Type I vs II | 1  2  3 | -6 -10 58  -34 -24 58  -22 -22 72 | 30  10  10 | 3.65  3.37  3.43 | Left  Left  Left | Medial Frontal Gyrus  Precentral Gyrus  Postcentral Gyrus |
| EPT | Mixed/  Item Type I  (Easy) | 1  2  3  4  5  6  7  8  9  10  11  12  13  14  15  16 | 2 12 42  28 -58 50  8 -88 -2  34 0 54  6 -72 50  -20 -32 6  -46 22 18  46 6 30  -10 0 6  -44 -68 -12  8 10 -2  2 -32 28  16 2 68  -44 34 22  -48 10 -2  14 -10 14 | 2717  731  602  237  189  109  56  51  30  23  23  22  18  15  15  12 | 11.2  13.4  7.81  5.73  9.80  4.69  5.50  6.28  4.35  5.23  4.84  3.65  3.74  4.05  5.23  3.88 | Left  Right  Right  Right  Right  Left  Left  Right  Left  Left  Right  Left  Right  Left  Left  Right | Medial Frontal Gyrus  Superior Parietal Lobule  Lingual Gyrus  Middle Frontal Gyrus  Precuneus  Thalamus  Middle Frontal Gyrus  Precentral Gyrus  Thalamus  Fusiform Gyrus  Caudate  Cingulate Gyrus  Superior Frontal Gyrus  Middle Frontal Gyrus  Insula  Thalamus |
|  | Mixed/  Item Type II (Hard) | 1  2  3  4  5  6  7  8  9  10  11  12  13  14 | 0 14 42  -48 2 42  28 -58 48  6 -72 50  -36 16 -6  40 16 -4  50 28 30  -48 10 0  0 -30 0  -34 48 26  -10 0 6  -6 -86 0  -40 -54 54  -4 -24 10 | 1760  1398  926  447  144  124  83  36  24  20  13  12  10  10 | 11.8  7.36  11.8  10.9  5.44  5.51  5.92  5.42  4.03  4.02  3.73  4.34  7.74  3.84 | Left Left  Right Right  Left  Right  Right  Left  Left  Left  Left  Left  Left  Left | Medial Frontal Gyrus  Precentral Gyrus  Superior Parietal Lobule  Precuneus  Insula  Insula  Precentral Gyrus  Insula  Thalamus  Middle Frontal Gyrus  Thalamus  Lingual Gyrus  Inferior Parietal Lobule  Thalamus |
| FT | Mixed/  Item Type I | 1  2  3  4  5  6  7  8  9  10  11  12 | -2 4 54  -46 2 30  12 -88 4  48 6 30  16 -16 16  22 -34 8  -42 12 -4  -46 -6 54  28 -58 64  -48 -66 -14  16 -4 18  30 -2 64 | 3077  1435  283  88  41  39  25  21  18  15  14  12 | 9.29  10.7  5.63  5.99  3.81  3.98  3.60  4.38  4.4  3.90  3.51  3.60 | Left  Left  Right  Right  Right  Right  Left  Left  Right  Left  Right  Right | Medial Frontal Gyrus  Precentral Gyrus  Lingual Gyrus  Precentral Gyrus  Thalamus  Thalamus  Insula  Precentral Gyrus  Superior Parietal Lobule  Fusiform Gyrus  Caudate  Middle Frontal Gyrus |
|  | Mixed/  Item Type II | 1  2  3  4  5  6  7  8  9  10  11  12 | -48 4 30  22 -66 48  6 -74 50  24 -34 10  -20 -34 8  46 6 30  16 -2 18  4 -32 26  -36 48 18  -10 -64 64  6 -28 12  14 -74 10 | 3212  649  225  137  126  70  54  53  50  28  27  14 | 13.2  9.71  8.71  5.51  4.39  6.31  4.28  4.36  4.42  6.37  4.12  3.42 | Left  Right  Right  Right  Left  Right  Right  Right  Left Left  Right  Right | Precentral Gyrus  Precuneus  Precuneus  Thalamus  Thalamus  Precentral Gyrus  Caudate  Posterior Cingulate  Middle Frontal Gyrus  Superior Parietal Gyrus  Thalamus  Lingual Gyrus |
| EPT | Mixed/  Item Type I vs II | 1  2  3  4  5  6  7 | -32 -26 66  -48 -6 2  0 -8 46  22 -58 4  2 12 26  44 -24 10  -44 -32 58 | 330  178  135  70  53  27  16 | 6.36  4.55  4.41  4.35  4.19  3.82  4.21 | Left  Left  Left  Right  Right  Right  Left | Postcentral Gyrus  Insula  Paracentral Lobule  Lingual Gyrus  Anterior Cingulate  Transverse Temporal Gyr.  Inferior Parietal Lobule |
|  | Mixed/  Item Type II vs I | 1  2  3  4  5  6 | -4 -60 38  -40 26 34  26 14 42  -24 16 38  -32 -2 30  -6 28 34 | 108  94  75  37  11  10 | 5.23  4.12  3.96  3.59  3.54  3.3 | Left  Left  Right  Left  Left  Left | Precuneus  Precentral Gyrus  Cingulate Gyrus  Middle Frontal Gyrus  Precentral Gyrus  Cingulate Gyrus |
| FT | Mixed/  Item Type I vs II | 1  2  3  4  5  6  7  8 | -2 -10 38  22 -24 74  -38 -30 56  -40 -20 56  -12 -32 36  52 -12 40  22 -50 2  14 -26 -6 | 1259  284  216  53  43  41  36  14 | 6.05  4.09  5.17  4.86  3.86  3.89  3.68  3.49 | Left  Right  Left  Left  Left  Right  Right  Right | Cingulate Gyrus  Precentral Gyrus  Inferior Parietal Gyrus  Postcentral Gyrus  Cingulate Gyrus  Precentral Gyrus  Parahippocampal Gyrus  Thalamus |
|  | Mixed/  Item Type II vs I | 1  2  3  4  5  6 | 6 -72 50  -28 10 50  28 14 48  -50 14 36  24 28 48  -36 24 32 | 312  244  97  41  25  12 | 5.22  4.49  4.10  3.83  3.91  3.90 | Right  Left  Right  Left  Right  Left | Precuneus  Middle Frontal Gyrus  Middle Frontal Gyrus  Middle Frontal Gyrus  Superior Frontal Gyrus  Middle Frontal Gyrus |

S3.0 Local activations by birth group.

The individual subject fMRI maps were investigated to characterize the individual hemodynamic responses during the experimental and baseline conditions. The MNI location of the peak voxel within each cluster was extracted using FSL’s ‘cluster’ command. A minimum cluster size of 10 voxels and a minimum t-score threshold of 3.1 (equivalent to p = 0.001) were implemented. The coordinates of the peak t-score within each cluster were converted to Talairach space using GingerALE software (<https://www.brainmap.org/ale/>) so that the anatomical locations could be determined using the Talairach Daemon (<http://www.talairach.org/daemon.html>). Each location was assigned a unique code from 1 to 105, corresponding to the full brain segmentation of the MNI template.

An exploratory Fisher’s exact test was performed to test the association between activation status in a range of locations with birth group. Counts of participants with activations within each region were compared by birth group. Two-sided significance level of 0.05 was adopted. Improper fractions, Item Type II (easier level) vs rest had a positive association with the EPT group for left parahippocampal gyrus (p = 0.035). Improper fractions, Item Type I (harder level) vs rest was associated with activity in the right medial frontal gyrus (p = 0.0131) and precuneus (p = 0.038). The mixed fractions, Item Type II (harder level) vs rest was associated with left medial frontal gyrus (p = 0.029) and superior temporal gyrus (p = 0.029) in the EPT group. No other significant associations were found across the other remaining contrasts.

S4.0 Linear SVM Classification

S4.1 Linear SVM methods

To further explore the activation differences between birth groups and strategy adoption, a linear support vector machine (SVM) analysis was performed. The aim was to determine if it was possible to systematically distinguish activation patterns in each of the item types in an automated way. Preprocessing of BOLD signal were performed using FEAT, as described above. Additional preprocessing steps via PyMVPA toolbox [7] included detrending, normalization, and ANOVA-based feature selection (using top 5% of active voxels) as described in [7]. Rest volumes were excluded from analysis. The linear SVM was implemented using a leave-one-out cross validation approach. For the group level analyses, a single subject’s mean activation for all conditions related to a specific strategy type (or strategy-item type combination) was used in classification to perform a leave-one-(subject)-out cross validation. As a result, the SVM would be trained on all strategy conditions from n-1 subjects prior to testing on the subject left out. Overall classification accuracies were determined using the PyMVPA toolbox built-in statistics.

S4.2 Linear SVM results

The purposes of these analyses were to determine if 1) birth group membership can be distinguished from hemodynamic response patterns from each strategy and item type, 2) difficulty level within a birth group and for a given strategy, and 3) strategy adoption within each birth group and for each of the item types. Discriminability with SVM corroborates findings of activation differences with the GLM-based analyses.

Classification of difficulty level (i.e., distinguishing item types) for the mixed fractions strategy demonstrated accuracies of 56.67% and 71.43% for FT and EPT groups, respectively (Figure 6a). Difficulty level classification for the improper fractions strategy demonstrated accuracies of 63.33% and 61.54% for FT and EPT groups, respectively. A similar analysis was performed to classify different strategies within the same item type. Classification of mixed versus improper strategies for Item Type II (improper easy vs mixed fractions hard) demonstrated accuracies of 86.67% and 80.77% for FT and EPT groups, respectively (Figure 6b). Strategy classification for Item Type I (improper hard vs mixed fractions easy) demonstrated accuracies of 70% and 69.23% for FT and EPT groups, respectively.

Figure 6

A)

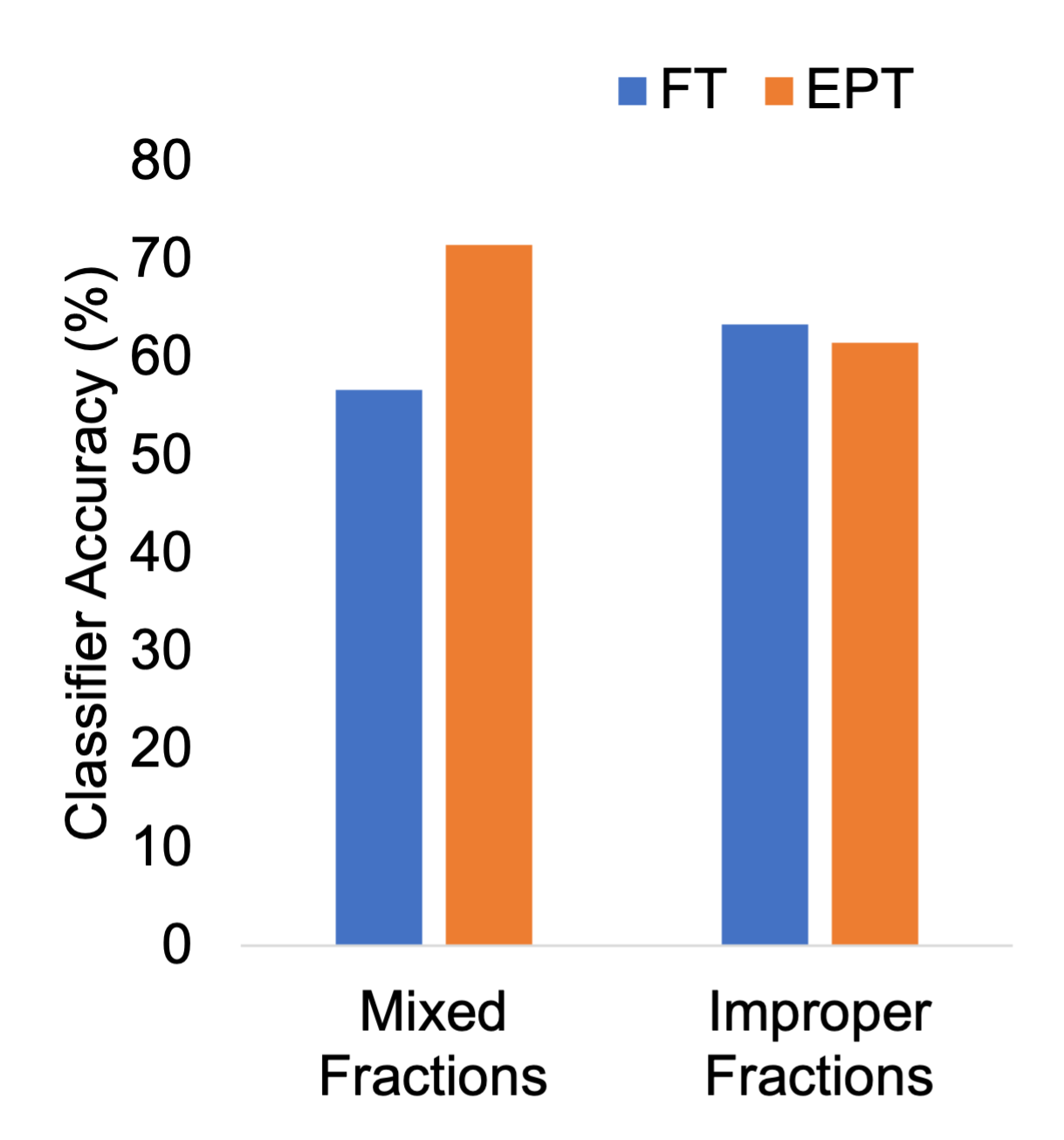

**B)**

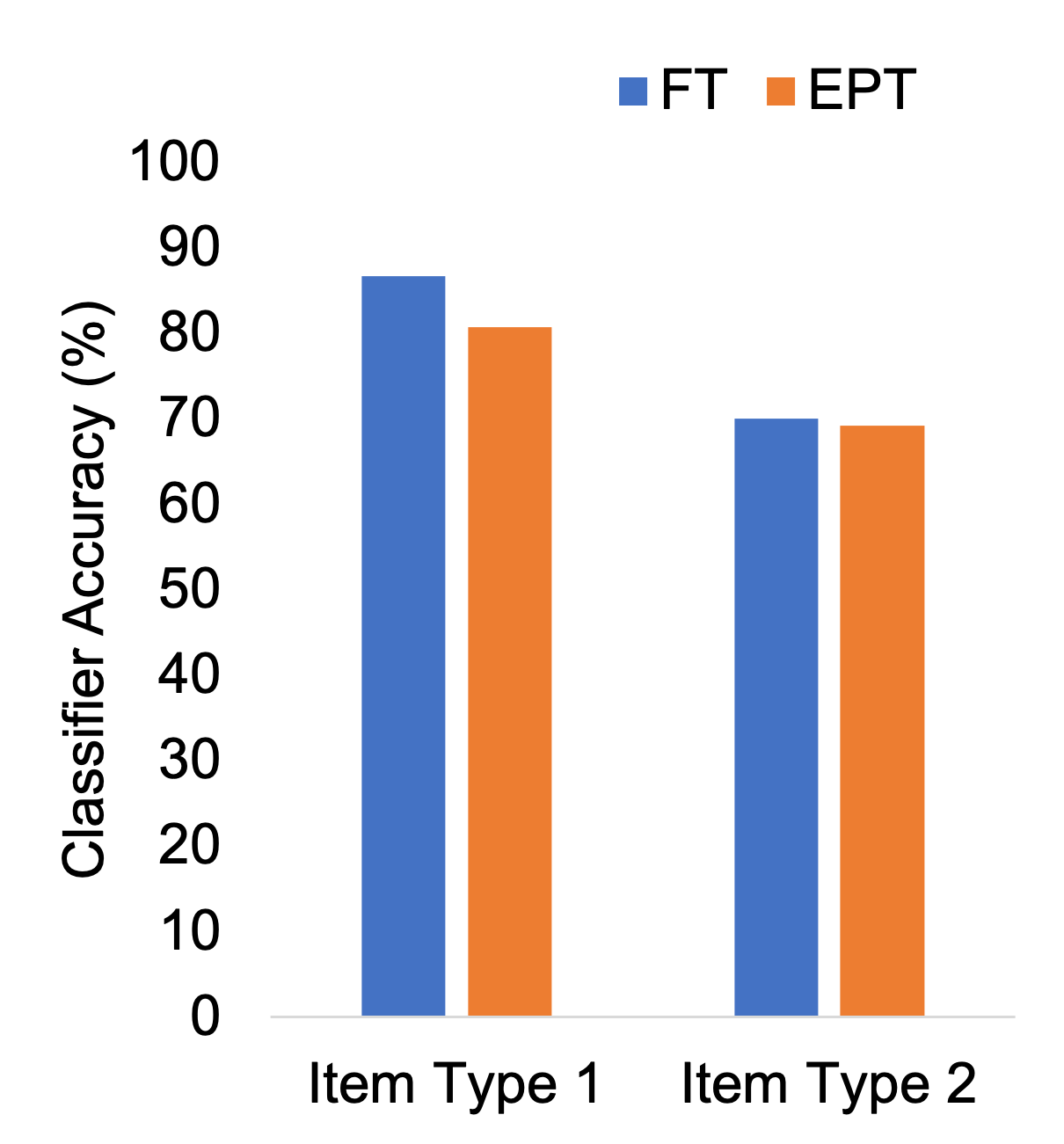

**Figure 6:** Accuracies of group-level SVM classification of difficulty level (A) and strategy (B) from both FT and EPT groups. A leave-one-out cross validation approach was used for all SVM analyses.

A further goal was to determine if the task-related BOLD signal measured here could be used by the SVM to identify specific traits about a subject, or more specifically, if the SVM could distinguish between birth status. BOLD signal data from a specific strategy and an item type (e.g., harder level of mixed fractions) was used with the goal of classifying untested BOLD signal data as belonging to an EPT versus FT subject. This was conducted over multiple cross validation iterations. The resulting classification accuracies and confusion matrices for each strategy-difficulty combination are detailed in Table 4. The strategy-difficulty combinations with the hardest difficulty produced the highest classification accuracies (67.86% and 64.29% for mixed and improper fractions, respectively).

Table 4

|  | **Classifier Accuracy (%)** | **Confusion Matrix (Prediction – Target)** | | | |
| --- | --- | --- | --- | --- | --- |
|  |  | **EPT-EPT** | **EPT-FT** | **FT-EPT** | **FT-FT** |
| Mixed, Item Type I (Easy) | 60.71 | 7 | 5 | 6 | 10 |
| Mixed, , Item Type II (Hard) | 67.86 | 6 | 2 | 7 | 13 |
| Improper, , Item Type II (Easy) | 28.57 | 1 | 8 | 12 | 7 |
| Improper, Item Type I (Hard) | 64.29 | 7 | 4 | 6 | 11 |
| Item Type I  (Mixed Easy, Improper Hard) | 53.57 | 6 | 6 | 7 | 9 |
| Item Type II  (Mixed Hard, Improper Easy) | 50.00 | 3 | 4 | 10 | 11 |

**Table 4.** Accuracies and confusion matrices from SVM classification of birth status.

S5.0 Volumetric analysis

S5.1 Volumetric analysis methods.

A volumetric analysis was performed on the T1 image for all subjects to investigate differences in grey and white matter volumes between the FT and EPT groups. T1 images were segmented using Freesurfer software (<https://freesurfer.net>) and were partitioned based on the Desikan-Killiany atlas [8]. This segmented whole-brain grey and white matter, producing 70 regions for each hemisphere and 6 subcortical grey matter regions (152 regions in total). The regions were evenly divided between the two hemispheres. Each region was coded with a unique number for easy identification and extraction for analysis. The command ‘fslmaths’ was used to threshold the outputted images to isolate each region of interest (ROI) and the total number of voxels within that ROI was obtained using ‘fslstats’. The ROI volume was normalized by total intracranial volume (obtained from Freesurfer output) to account for differences in brain sizes across all the subjects. A two-tailed t-test was run for each ROI comparing the normalized volumes in the EPT and FT groups. The result was considered significant if α ≤ 0.05. Cohen’s D values were also calculated to give an indication of magnitude of difference between the groups. A Cohen’s D value ≥ |0.8| was considered significant and a negative value indicated increased volume in the EPT group while a positive value indicated decreased volume.

S5.2 Volumetric results.

The EPT group compared to the FT group showed decreased grey matter volume in left lingual and parahippocampal gyri, superior and middle temporal gyri, left pallidum and bilateral thalami. Increased grey matter volume in EPT relative to FT participants was found in right inferior temporal gyrus.

The EPT group compared to the FT group showed decreased white matter volume in left parahippocampal and middle frontal gyri, posterior cingulate; right superior temporal sulcus; and bilateral lingual gyri and superior parietal lobules. Increased white matter volume in the EPT group was also observed in bilateral middle temporal gyri. All volumetric differences are given in Table S2.

**Table S2:** Grey and white matter volume differences between the EPT and FT groups. α ≤ 0.05 and Cohen’s D value ≥ 0.8.

| **Hemisphere** | **Matter** | **Region** | **α** | **Cohen’s D** |
| --- | --- | --- | --- | --- |
| **Decreased** |  |  |  |  |
| Left | Grey | Lingual gyrus | 0.0009 | 1.1306 |
| Left | Grey | Parahippocampal gyrus | 0.0138 | 0.8766 |
| Right | Grey | Banks of the Superior Temporal sulcus | 0.0030 | 1.0304 |
| Right | Grey | Middle temporal gyrus | 0.0078 | 0.9393 |
| Decreased |  |  |  |  |
| Left | Subcortical | Pallidum | 0.0164 | 0.8567 |
| Left | Subcortical | Thalamus | 0.0238 | 0.8115 |
| Right | Subcortical | Thalamus | 0.0207 | 0.8286 |
| **Increased** |  |  |  |  |
| Right | Grey | Inferior Temporal gyrus | 0.0066 | -0.9556 |
| **Decreased** |  |  |  |  |
| Left | White | Lingual gyrus | 0.0044 | 0.9960 |
| Left | White | Parahippocampal gyrus | 0.0188 | 0.8406 |
| Left | White | Caudal Middle frontal gyrus | 0.0091 | 0.9227 |
| Left | White | Isthmus Cingulate (posterior area) | 0.0015 | 1.0896 |
| Left | White | Superior Parietal lobule | 0.0084 | 0.9304 |
| Right | White | Banks of the Superior Temporal sulcus | 0.0092 | 0.9212 |
| Right | White | Lingual gyrus | 0.0058 | 0.9681 |
| Right | White | Superior Parietal lobule | 0.0076 | 0.9409 |
| **Increased** |  |  |  |  |
| Left | White | Middle temporal gyrus | 0.0012 | -1.1101 |
| Right | White | Middle temporal gyrus | 0.0038 | -1.0088 |

S6.0 DTI Analysis

S6.1 DTI methods.

Based on individual-level tractography, we analyzed and compared structural connectivity between EPT and control groups, as described in [9]. Wavelet representations of connectivity graphs were derived, with streamline counts data used to reflect connectivity between ROIs. Multivariate analysis methods applied to wavelet coefficients allow for efficient joint testing of group differences between specific ROIs.

The diffusion weighted images were pre-processed using both FSL and MRtrix (www.mrtrix.org). Data were preprocessed by performing denoising, motion correction and eddy current correction. A whole brain mask was applied to remove voxels outside the brain before estimating the response functions for spherical deconvolution using the Tournier algorithm in MRtrix. Finally, the fiber orientation distribution was estimated using constrained spherical deconvolution [10]. The T1 images were transformed to DTI space and brain extracted. Segmentation of grey, white matter, cerebrospinal fluid and subcortical structures was performed and a composite image was composed to inform and anatomically constrain the DTI tractography [11]. Whole brain probabilistic tractography was performed in MRtrix using second order integration over the fiber orientations distributions (iFOD2 algorithm) [12].

Areas of the brain relevant to the mathematical network were identified from a literature survey. These included: frontal eye fields, middle and inferior frontal gyri, supramarginal gyrus, inferior and superior parietal lobules, lingual gyrus, parahippocampal gyrus, insula and visual area (cuneus and lateral occipital cortex). They were extracted from the individual subject Freesurfer segmented image (see Section: “Volumetric Analysis”). Each extracted region of interest was binarized and multiplied by a unique number from 1-22 and then all were added together to form a single image. The purpose was to create a unique image intensity value for each ROI so they could be distinguished in the final single image. The final ROI image was then applied to the resulting tracks to determine the number of connections that existed between each ROI and every other ROI. These values were adjusted for differences in intracranial volume [13, 14]. A weighting was also applied using the average FA and mean diffusivity (MD) along the streamline tracks so that the final connectivity values were derived by:

$$streamline count \cdot mean\_FA \cdot mean\_MD \cdot\frac{(volume of ROI 1+volume of ROI 2)}{intracranial volume}$$

There was no directionality to the connections – i.e. an area 1 to area 2 connection is identical to an area 2 to area 1 connection.

The multiresolution brain connectivity toolbox (MBCA, [9, 15]), run through Matlab ([www.mathworks.com](http://www.mathworks.com)) performed the connectivity analysis using wavelets. Methods are described in detail in [9] and [15]. Connectivity changes were considered significant if they were below the threshold p < 0.05.

S6.2 DTI results.

An exploratory analysis of differences between birth groups in structural brain connectivity showed a number of areas where the EPT group displayed decreased connectivity compared the FT group, while only two areas displayed increased connectivity. The results are presented in Figure S4 for both increased and decreased connectivity. Weaker connectivity in the EPT compared to the FT group was observed between the left visual area and the left inferior parietal lobule and right fusiform and lingual gyri; between the left lingual gyrus and the left insula and right parahippocampal gyrus; between the left parahippocampal gyrus and the left superior parietal lobule and right lingual and parahippocampal gyris and superior parietal lobule; between the left superior parietal lobule and the left supramarginal gyrus; between the right visual area and the right superior parietal lobule; between the right fusiform gyrus and the right parahippocampal gyrus; between the right lingual gyrus and the right insula; and between the right superior parietal lobule and the right supramarginal gyrus. Increased connectivity in the EPT compared to the FT group was observed between the left parahippocampal gyrus and right fusiform gyrus, and between the right superior parietal lobule and the right insula.

**Figure S4:** A connectivity matrix showing connections that differed between the EPT and the FT group. Regions are listed on each axis. Connectivity was considered significant if p < 0.05.

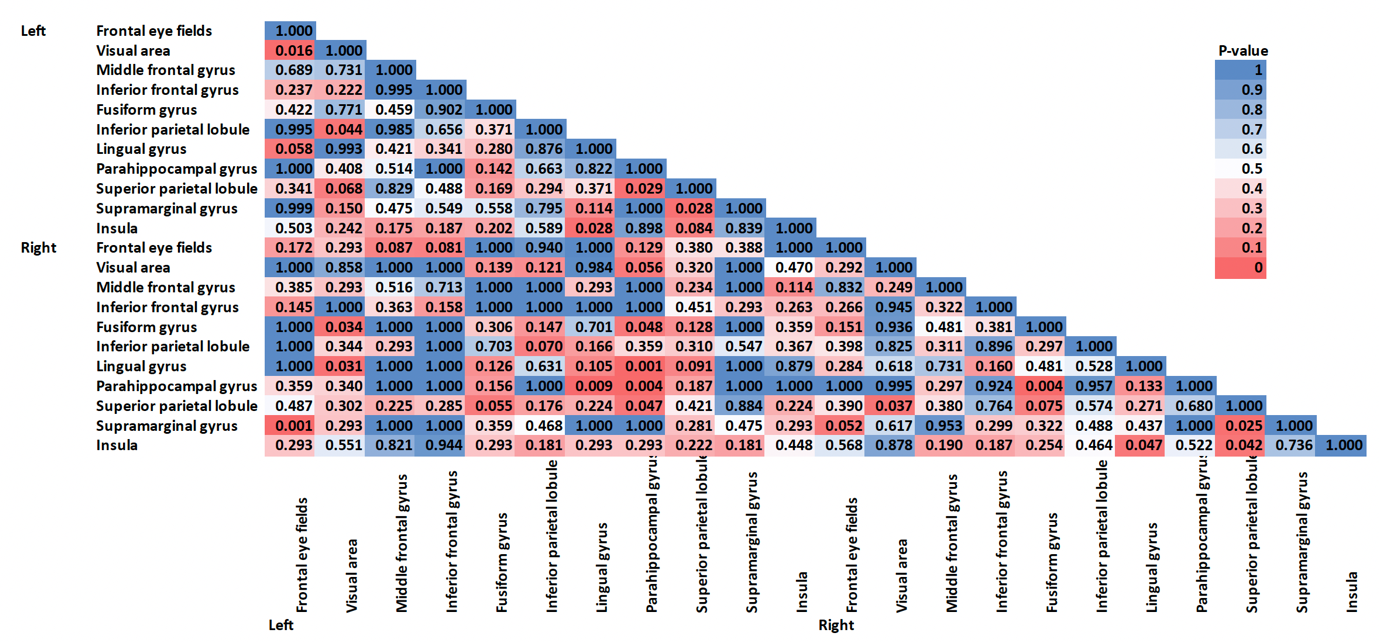
